# Astrocytic striatal GABA transporter activity governs dopamine release and shows maladaptive downregulation in early parkinsonism

**DOI:** 10.1101/698274

**Authors:** Bradley M. Roberts, Natalie M. Doig, Katherine R. Brimblecombe, Emanuel F. Lopes, Ruth E. Siddorn, Sarah Threlfell, Natalie Connor-Robson, Nora Bengoa-Vergniory, Nicholas Pasternack, Richard Wade-Martins, Peter J. Magill, Stephanie J. Cragg

## Abstract

Striatal dopamine (DA) is critical for action and learning. Recent data show DA release is under tonic inhibition by striatal GABA. Ambient striatal GABA tone on striatal projection neurons can be governed by plasma membrane GABA uptake transporters (GATs) on astrocytes. However, whether striatal GATs and astrocytes determine DA output are unknown. We reveal that DA release in mouse dorsolateral striatum, but not nucleus accumbens core, is governed by GAT-1 and GAT-3. These GATs are partly localized to astrocytes, and are enriched in dorsolateral striatum compared to accumbens core. In a mouse model of early parkinsonism, GATs were downregulated and tonic GABAergic inhibition of DA release augmented, with corresponding attenuation of GABA co-release from dopaminergic axons. These data define previously unappreciated and important roles for GATs and astrocytes in determining DA release in striatum, and reveal a maladaptive plasticity in early parkinsonism that impairs DA output in vulnerable striatal regions.

**Highlights:**

1. GABA transporters set the level of GABA inhibition of DA output in dorsal striatum
2. Astrocytes facilitate DA release levels by limiting tonic GABA inhibition
3. Tonic inhibition of DA release is augmented in a mouse model of early parkinsonism
4. DA and GABA co-release are reduced in a mouse model of early parkinsonism

## INTRODUCTION

Dopamine (DA) release in the dorsal and ventral striatum plays key roles in action selection and motivation, and is dysregulated in diverse disorders including Parkinson’s disease (PD) and addictions. Striatal DA release is gated locally by axonal mechanisms and striatal neuromodulators that regulate or even drive DA release (Schmitz et al., 2003; Sulzer et al., 2016). It has recently been revealed that DA release is under tonic inhibition by striatal GABA, operating through GABA_A_ and GABA_B_ receptors (Lopes et al., 2019; Pitman et al., 2014; Schmitz et al., 2002). The striatum contains a high density of GABAergic projection neurons and interneurons and also receives a source of GABA co-released from mesostriatal DA neurons (Kim et al., 2015; Tritsch et al., 2012, 2014). Given the paucity of GABAergic synapses on DA axons (Charara et al., 1999), tonic inhibition of DA release by striatal GABA is presumably mediated through extrasynaptic effects of ambient GABA (Lopeset al., 2019) on receptors located presumably on DA axons. GABA can spill over for extrasynaptic actions in other nuclei (Farrant and Nusser, 2005), and in the dorsal striatum, provides a sizeable ambient GABA tone on spiny projection neurons (SPNs), evident as a tonic GABA_A_ receptor-mediated inhibitory conductance (Ade et al., 2008; Cepeda et al., 2013; Kirmse et al., 2008, 2009; Santhakumar et al., 2010).

Tonic inhibition by ambient GABA across the mammalian brain is usually limited by uptake by plasma membrane GABA transporters (GATs) (Brickley and Mody, 2012). There are two isoforms of the GAT in striatum: GAT-1 (*Slc6a1*), abundant in axons of GABAergic neurons (Augood et al., 1995; Durkin et al., 1995; Ng et al., 2000; Yasumi et al., 1997); and GAT-3 (*Slc6a11*), expressed moderately (Ficková et al., 1999; Ng et al., 2000; Yasumi et al., 1997) but observed particularly on astrocytes (Chai et al., 2017; Ng et al., 2000; Yu et al., 2018). Emerging transcriptomic data additionally indicate that striatal astrocytes express both GAT-3 and GAT-1 (Chai et al., 2017; Gokce et al., 2016; Zhang et al., 2014). In addition, mRNA for GAT-1 and for GAT-3 has been found in midbrain DA neurons and these GATs have been suggested but not confirmed to be located on striatal DA axons to support GABA co-storage and co-release (Tritsch et al., 2014). Ambient GABA tone on SPNs in dorsal striatum is limited by the activity of GAT-1 and GAT-3 (Kirmse et al., 2008, 2009; Santhakumar et al., 2010; Wójtowicz et al., 2013), and recent evidence indicates that dysregulation of GAT-3 on striatal astrocytes results in profound changes to SPN activity and striatal-dependent behavior (Yu et al., 2018). However, whether striatal GAT function and, by association, astrocytes are critical for setting the level of DA output has not previously been examined.

Here we reveal that GAT-1 and GAT-3 strongly regulate striatal DA release in the dorsolateral striatum (DLS) but not in the nucleus accumbens core (NAcC), by limiting tonic inhibition arising from striatal ambient GABA. We identify a role for GATs located on striatal astrocytes in supporting DA release, and furthermore, reveal maladaptive reductions in GAT levels that impair DA output in the DLS in a mouse model of early parkinsonism.

## RESULTS

### DA release in DLS and NAcC is tonically inhibited by a GAD-dependent GABA source

We recently reported that axonal DA release in the dorsal striatum is under tonic inhibition by striatal GABA, as GABA_A_ and GABA_B_ receptor antagonists enhanced DA release evoked by single electrical and targeted optogenetic stimuli (Lopes et al., 2019). Since mechanisms that regulate striatal DA release can diverge between dorsal and ventral striatal territories (Brimblecombe et al., 2015; Britt and McGehee, 2008; Janezic et al., 2013; Shin et al., 2017; Threlfell and Cragg, 2011; Threlfell et al., 2010), we first determined whether DA release in NAcC, within the ventral striatum, is similarly regulated by tonic GABA inhibition. We used fast-scan cyclic voltammetry (FSCV) in acute coronal slices of mouse brain to detect extracellular concentration of DA ([DA]_o_) at carbon-fiber microelectrodes evoked optogenetically to activate DA axons selectively (**Fig. 1a-b**). Co-application of GABA_A_ and GABA_B_ receptor antagonists (+)-bicuculline (10 μM) and CGP 55845 (4 μM) respectively, significantly enhanced [DA]_o_ evoked by single light pulses by ∼25% in both DLS and NAcC, when compared to time-matched drug-free controls (**Fig. 1c-d**; DLS: F_(1,16)_ = 33.92, p < 0.0001; two-way repeated-measures; time-matched controls: n = 8 experiments/5 mice, GABA_R_ antagonists: n = 10 experiments/5 mice; NAcC: F_(1,12)_ = 20.68, p = 0.0007; two-way repeated-measures ANOVA; time-matched controls: n = 6 experiments/5 mice, GABA_R_ antagonists: n = 8 experiments/5 mice). These effects were similar in DLS and NAcC (**Fig. 1e**; t_(16)_ = 1.089, p = 0.292; unpaired Student’s t-test) and were also observed when [DA]_o_ was evoked by single electrical pulses (**Supplementary Fig. 1a-c**). This tonic GABAergic inhibition of DA release did not require cholinergic interneuron input to nAChRs (**Supplementary Fig. 1**), striatal glutamatergic input (**Supplementary Fig. 1d**), and was also seen when higher near-physiological bath temperatures of 37 °C were used (**Supplementary Fig. 1e**). These results confirm that DA release is under tonic inhibition by GABA in both ventral and dorsal striatal regions.

**Fig 1.**
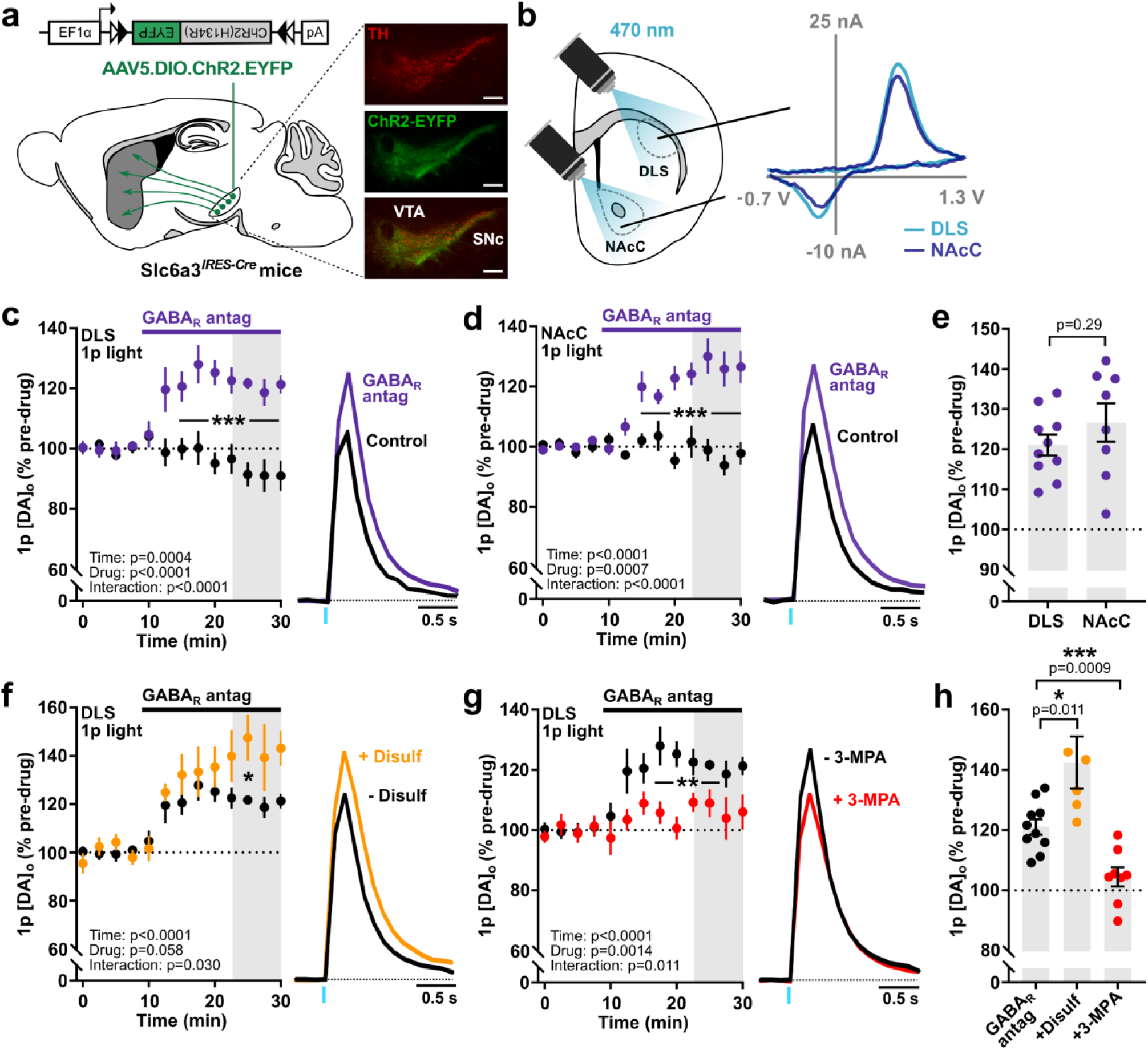
Striatal DA release is tonically inhibited by a GAD-dependent GABA source. (**a, b**) Schematics representing the experimental configuration and representative voltammograms for light-evoked [DA]_o_ in DLS and NAcC, after injection and expression of viral ChR2-eYFP in VTA and SNc in Slc6a3^*IRES-Cre*^ mouse. TH (*red*), ChR2-eYFP (*green*). Scale bars: 0.25 mm. (**c, d**) *Left*, mean peak [DA]_o_ (± SEM) during consecutive recordings evoked by a light pulse (*1p*) in control conditions (*black*, n = 8 experiments/6 mice for DLS, n = 6 experiments/5 mice for NAcC) and with GABA_A_ and GABA_B_ receptor antagonists (*solid bar*)(*purple, GABA*_*R*_ *antag*), (+)-bicuculline (10 μM) and CGP 55845 (4 μM), respectively, recorded in the DLS (c, n = 9 experiments/5 mice) or NAcC (d, n = 5 experiments/4 mice). *Right*, mean transients of [DA]_o_ (normalized to pre-drug baselines) from last 4 time points (gray shaded region). (**e**) Mean peak [DA]_o_ (± SEM) evoked by 1p following GABA_R_ antagonism in DLS and NAcC (as % of pre-drug baseline). (**f, g**) Mean peak [DA]_o_ (± SEM) during consecutive recordings evoked by 1p light during application of GABA_R_ antagonists in the absence (*black*, n = 10 experiments/7 mice) or the presence of ALDH inhibitor disulfiram (10 μM) (f, *orange*, n = 6 experiments/5 mice) or GAD inhibitor 3-MPA (5000 μM) (g, *red*, n = 7 experiments/5 mice). (**h**) Mean peak [DA]_o_ (± SEM) in DLS following GABA_R_ antagonism in the absence or the presence of 3-MPA and disulfiram (as a % of pre-drug baseline). Data are normalized to mean of 4 time points prior to GABA antagonist application (*dotted line*).Mean transients of [DA]_o_ are derived from last 4 time points (*gray shaded region*) and normalized to pre-drug baselines. Two-way repeated-measures ANOVA with Sidak’s multiple comparison tests (c, d, f, g) and Student’s unpaired t-tests (e, h). \**p* < 0.05, \*\**p* < 0.01, \*\*\**p* < 0.0001.

We tested whether GABAergic inhibition of DA release arose from GABA co-released by DA axons or from GABA originating from a canonical neuron source (i.e. striatal GABAergic neurons). Mesostriatal DA neurons synthesize, co-store and co-release GABA (Tritsch et al., 2012), with GABA synthesis depending on aldehyde dehydrogenase (ALDH)-1a1 (Kim et al., 2015). In contrast, canonical synthesis of GABA in neurons requires glutamate decarboxylase (GAD). We examined which source(s) of GABA is responsible for tonic inhibition of DA release using inhibitors of either ALDH or GAD. Pre-treating slices with a non-selective ALDH inhibitor disulfiram (10 μM, 2-4 hrs) halved light-evoked GABA_A_-dependent currents from DA axons onto SPNs (**Supplementary Fig. 2a**), as reported previously (Kim et al., 2015), but did not prevent GABA receptor antagonists from enhancing DA release: in the DLS, GABA receptor antagonists enhanced light-evoked [DA]_o_ by ∼40% in the presence of disulfiram, which was a significantly larger effect than seen without disulfiram (**Fig. 1f**; F_(12,168)_ = 1.97, p = 0.030; two-way repeated-measures ANOVA, drug x time interaction; **Fig. 1h**; t_(14)_ = 2.923, p = 0.011; unpaired Student’s t-test; disulfiram: n = 6 experiments/5 mice, disulfiram absent: n = 10 experiments/5 mice). Disulfiram alone did not significantly modify evoked [DA]_o_ (**Supplementary Fig. 2b**). These data suggest that GABA co-released from DA axons is not responsible for tonic inhibition of DA release, and rather, that an ALDH-dependent source of GABA might act indirectly to limit tonic inhibition of DA by a different, ALDH-independent source. By contrast, when we pre-treated brain slices with the GAD inhibitor 3-mercaptopropionic acid (3-MPA, 500 μM), which attenuates electrically evoked GABA transmission onto SPNs by more than half (Kim et al., 2015), the disinhibition of DA release in the DLS by GABA receptor antagonists was attenuated (**Fig. 1g**; F_(1,16)_ = 14.81, p = 0.0014; two-way repeated-measures ANOVA; **Fig. 1h**; t_(16)_ = 4.056, p = 0.0009; unpaired Student’s t-test; 3-MPA: n = 8 experiments/5 mice, 3-MPA absent: n = 10 experiments/5 mice), indicating that a GAD-dependent GABA source provides tonic inhibition of striatal DA release.

### GAT-1 and GAT-3 inhibition attenuates DA release in the DLS but not NAcC

We tested the hypothesis that GATs, by governing ambient GABA (Kirmse et al., 2008, 2009; Santhakumar et al., 2010; Wójtowicz et al., 2013), might determine the level of tonic inhibition of DA release. The non-selective GAT inhibitor (±)-nipecotic acid (NPA) (1 - 10 mM) inhibits all subtypes of GATs (Goubard et al., 2011; Li et al., 2017). Bath application of NPA (1.5 mM) attenuated [DA]_o_ by single electrical pulses in the DLS to ∼60% of time-matched controls (**Fig. 2a**; F_(1,16)_ = 73.40, p < 0.0001; two-way repeated-measures ANOVA; NPA: n = 9 experiments/5 mice, time-matched controls: n = 9 experiments/7 mice). NPA also significantly reduced [DA]_o_ evoked optogenetically by single light pulses (**Supplementary Fig. 3a**), indicating that attenuation of DA release does not require concurrent activation of other striatal neurons. NPA attenuated electrically evoked [DA]_o_ to a greater extent in DLS than in NAcC (**Fig. 2a-c**; t_(13)_ = 5.266, p = 0.0002; unpaired Student’s t-test) where NPA only marginally attenuated DA release compared to time-matched controls (**Fig. 2b**; F_(1,10)_ = 6.72, p = 0.027; two-way repeated-measures ANOVA; NPA: n = 6 experiments/4 mice, time-matched controls: n = 6 experiments/5 mice). These data indicate that the level of tonic inhibition of DA release is limited by GATs in DLS to a greater degree than in NAcC.

**Fig 2.**
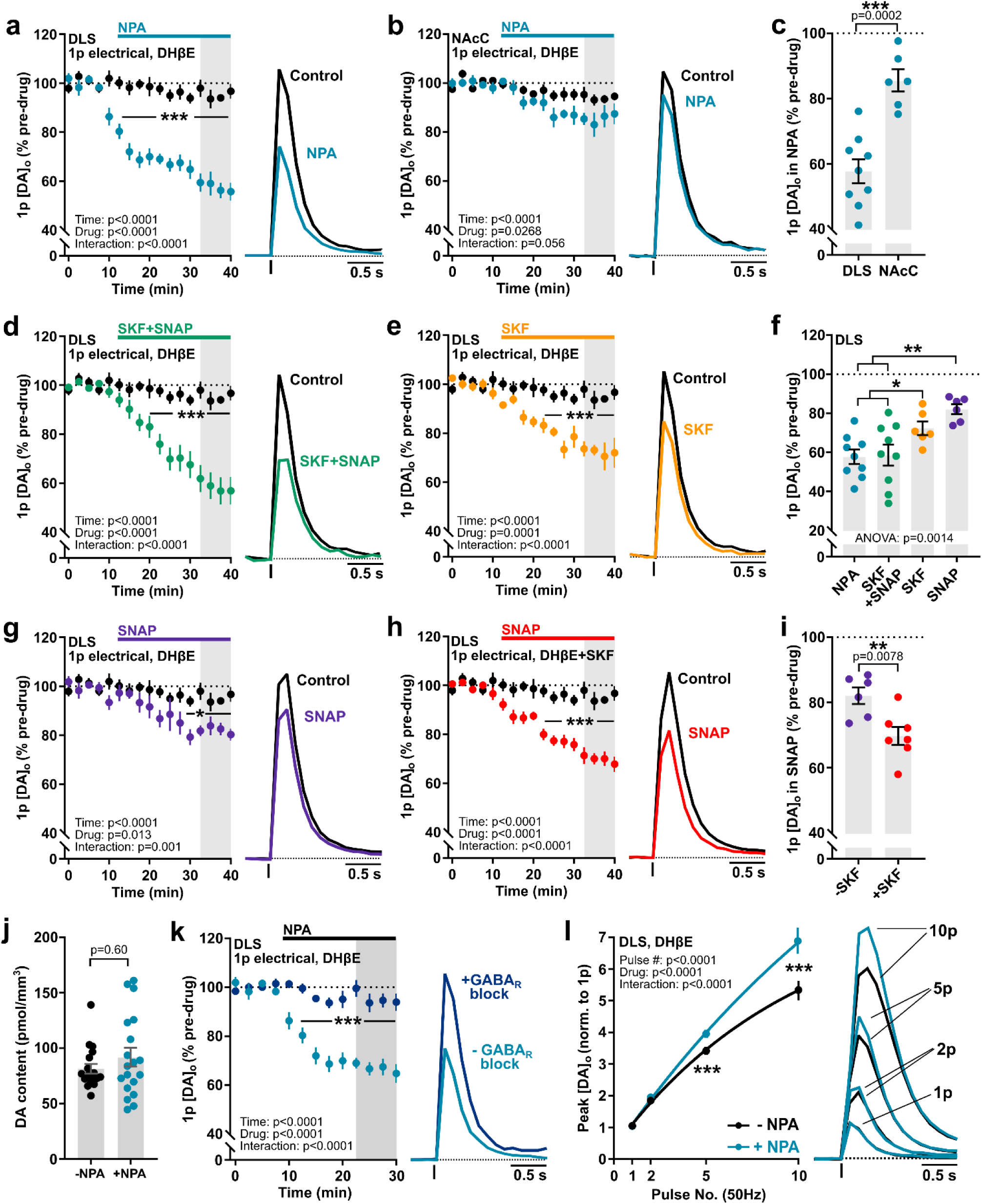
GAT-1 and GAT-3 inhibition attenuates DA release in DLS, but not NAcC. (**a**-**b, d**-**e, g**-**h**) Mean peak [DA]_o_ (± SEM) during consecutive recordings evoked by a single electrical pulse in DLS (a, d-e, g-h) or NAcC (b) in control conditions (*black*, n = 9 experiments/7 mice for DLS, n = 6 experiments/5 mice for NAcC) or with GAT inhibitor nipecotic acid (NPA, 1.5 mM) (a, *blue*, n = 9 experiments/5 mice; b, n = 6 experiments/4 mice), combined bath application of the GAT-1 specific inhibitor SKF89976A (20 μM) and the GAT-3 specific inhibitor SNAP5114 (50 μM) (d, *green*, n = 9 experiments/5 mice), SKF89976A alone (e, *orange*, n = 6 experiments/4 mice), SNAP5114 alone (g, *purple*, n = 6 experiments/4 mice), or bath application of SNAP5114 in slices pre-incubated in SKF89976A (h, *red*, n = 7 experiments/5 mice). (**c**, **f, i**) Mean peak [DA]_o_ (± SEM) evoked by 1p following GAT inhibition (expressed as a % of pre-drug baseline). (**j**) Mean DA content (± SEM) of dorsal striatum incubated in vehicle-treated control conditions (*black*, n = 19 punches/5 mice) or NPA (1.5 mM) (*blue*, n = 19 punches/5 mice). (**k**) Mean peak [DA]_o_ (± SEM) during consecutive recordings evoked by a 1 electrical pulse in DLS during application of NPA (1.5 mM) in the absence (*dark blue*, n = 9 experiments/5 mice) or presence (*light blue*, n = 5 experiments/4 mice) of GABA_A_ (picrotoxin, 100 μM) and GABA_B_ (CGP 55845, 4 μM) receptor antagonists. (**l**) *Left*, Mean peak values of [DA]_o_ evoked by 50 Hz electrical pulses in DLS normalized to 1p in the absence (*black*, control, n = 8 experiments/5 mice) or presence of NPA (1.5 mM) (*blue*, n = 8 experiments/5 mice). Sigmoidal curve fits (R^2^ = 0.98). Data are normalized to mean of 4 time points prior to GAT inhibitor application (*dotted line*); mean transients of [DA]_o_ are derived from last 4 time points (*gray shaded region*) and normalized to pre-drug baselines. DHβE (1 μM) present throughout. Two-way repeated-measures ANOVA with Sidak’s multiple comparison tests (a-b, d-e, g-h, k-l), Student’s unpaired t-tests (c, i), Mann-Whitney test (j) and one-way ANOVA with Sidak’s multiple comparison tests (f). \**p* < 0.05, \*\**p* < 0.01, \*\*\**p* < 0.0001.

Two main isoforms of GATs are expressed in the basal ganglia: GAT-1 and GAT-3 (Jin et al., 2011). We used selective inhibitors of GAT-1 and GAT-3 to identify which isoform(s) limit GABAergic inhibition of DA release in the DLS. Together, the combined inhibition of GAT-1 and GAT-3, with selective inhibitors SKF89976A (20 μM) and SNAP5114 (50 μM) respectively, significantly attenuated electrically evoked [DA]_o_ to ∼60% of time-matched controls (**Fig. 2d**; F_(1,16)_ = 24.79, p < 0.0001; two-way repeated-measures ANOVA; SKF+SNAP: n = 9 experiments/5 mice), equivalent to that seen with broad-spectrum GAT inhibitor NPA (**Fig. 2f**; F_(3,26)_ = 6.912, p = 0.0014, one-way ANOVA; SKF+SNAP vs. NPA, p = 0.9984, Sidak’s multiple comparisons). GAT-1 inhibition alone with SKF89976A (20 μM) significantly attenuated evoked [DA]_o_ in DLS to ∼75% of time-matched controls (**Fig. 2e**; F_(1,13)_ = 28.37, p = 0.0001; two-way repeated-measures ANOVA; SKF: n = 6 experiments/4 mice) and GAT-3 inhibition alone with SNAP5114 (50 μM) significantly attenuated evoked [DA]_o_ in DLS to ∼80% of time-matched controls (**Fig. 2g**; F_(1,13)_ = 8.205, p = 0.0133; two-way repeated-measures ANOVA; SNAP: n = 6 experiments/4 mice), which were smaller effects compared to combined GAT-1 and GAT-3 inhibition (**Fig. 2f**; F_(3,26)_ = 6.912, p = 0.0014, one-way ANOVA; SKF vs. NPA, p = 0.0192; SKF vs. SKF+SNAP, p = 0.0487; SNAP vs. NPA, p = 0.0031; SNAP vs. SKF+SNAP, p = 0.0045; Sidak’s multiple comparisons). The functional effects of GAT-3 inhibition on GABA_A_ receptor-mediated tonic currents in SPNs have been shown to be compensated for by GAT-1-mediated GABA uptake, with GAT-3 function revealed better during GAT-1 inhibition (Jin et al., 2011; Kirmse et al., 2009). We pre-treated slices with GAT-1 inhibitor SKF89976A (20 μM) and revealed that subsequent bath application of GAT-3 inhibitor SNAP5114 (50 μM) attenuated electrically evoked [DA]_o_ in DLS to ∼70% of time-matched controls (**Fig. 2h**; F_(1,14)_ = 35.61, p < 0.0001; two-way repeated-measures ANOVA; SNAP with SKF pre-treatment: n = 7 experiments/5 mice), a larger effect than with GAT-3 inhibitor alone (**Fig. 2i**; t_(11)_ = 3.243, p = 0.0078; unpaired Student’s t-test). Altogether, these data show roles for both GAT-1 and GAT-3 in limiting the level of GABA inhibition of DA release in the DLS.

### GAT inhibition attenuates striatal DA release by increasing GABA receptor tone

We ruled out diminished DA storage as a cause of the attenuation of DA release following GAT inhibition: Striatal DA content measured using high performance liquid chromatography (HPLC) with electrochemical detection was unchanged by incubation with GAT inhibitor NPA (**Fig. 2j**; U = 162, p = 0.603, Mann-Whitney test, n = 19 experiments/5 mice per condition). Instead, we confirmed that GAT inhibition modified DA release in a GABA-receptor dependent manner. The acute effects of NPA on evoked [DA]_o_ were prevented in the presence of antagonists for GABA_A_ (picrotoxin, 100 μM) and GABA_B_ (CGP55845, 4 μM) receptors (**Fig. 2k**; F_(1,12)_ = 40.41, p < 0.0001; two-way repeated-measures ANOVA; without GABA receptor antagonists: ∼65% of baseline, n = 9 experiments/7 mice; with GABA receptor antagonists: ∼95% of baseline, n = 5 experiments/4 mice), consistent with GAT regulation of DA release being mediated via extracellular GABA acting on GABA receptors. In addition, we excluded roles in the effects of NPA of D_2_ dopamine receptors, glutamate receptors, or modulation of DA uptake, since NPA effects were preserved in the presence of respective inhibitors of each of these potential mechanisms (**Supplementary Fig. 3b**,**c**). We have previously shown that activation of striatal GABA receptors can slightly promote the activity-dependence of DA release during short stimulus trains (Lopes et al., 2019). Consistent with an increase in GABA receptor activation, GAT inhibitor NPA increased the dependence of [DA]_o_ on pulse number during 50 Hz pulse trains in DLS (**Fig. 2l**; F_(1,7)_ = 128.9, p < 0.0001; two-way repeated-measures ANOVA; n = 8 experiments/5 mice). NPA also increased the paired-pulse ratio of electrically evoked [DA]_o_ at short inter-pulse intervals (**Supplementary Fig. 3d-e**) consistent with a decrease in DA release probability (Jennings et al., 2015). Together these data indicate that GAT inhibition attenuates DA release through increasing GABA receptor tone and reducing DA release probability.

We assessed whether the greater role for GATs in regulating [DA]_o_ in DLS than NAcC (see Fig. 2c) was due to differences in GABA receptor regulation of DA. However, bath application of exogenous GABA (2 mM) attenuated [DA]_o_ evoked by 1p electrical stimulation to a similar degree in DLS and NAcC (**Supplementary Fig. 3f**), arguing against a difference in GABA receptor function as a major factor. These findings therefore suggest a different level of GAT function in limiting ambient GABA in DLS versus NAcC.

### GAT-1 and GAT-3 function and expression is enriched in DLS versus NAcC

To identify whether GAT plays a greater role in governing GABA tone in DLS than NAcC, we recorded the tonic GABA_A_ receptor-mediated currents in SPNs using whole-cell voltage-clamp electrophysiology and assessed the impact of GAT inhibition on holding current. We confirmed that changes in holding current were mediated by GABA_A_ receptors by subsequently applying GABA_A_ receptor antagonist picrotoxin (PTX; 100 μM). Consistent with the differential effects on DA release, GAT inhibition with NPA (1.5 mM) promoted the GABA_A_-mediated holding current in SPNs to a greater degree in DLS than in NAcC (**Fig. 3a-c**; DLS: p = 0.0003, Friedman’s ANOVA on Ranks, NPA vs. drug-free baseline: p = 0.001, NPA + PTX vs. drug-free baseline: p = 0.16, NPA vs. NPA + PTX: p < 0.001, Student-Newman-Keuls tests, n = 7 cells/5 mice; NAcC: p = 0.0001, Friedman’s ANOVA on Ranks, NPA vs. drug-free baseline: p = 0.014, NPA + PTX vs. drug-free baseline: p = 0.014, NPA vs. NPA + PTX: p = 0.002, Student-Newman-Keuls tests, n = 6 cells/3 mice; DLS vs NAcC: U = 4, p = 0.0140, Mann-Whitney test). These data corroborate a greater role for GATs in limiting ambient GABA tone in DLS than in NAcC.

**Fig 3.**
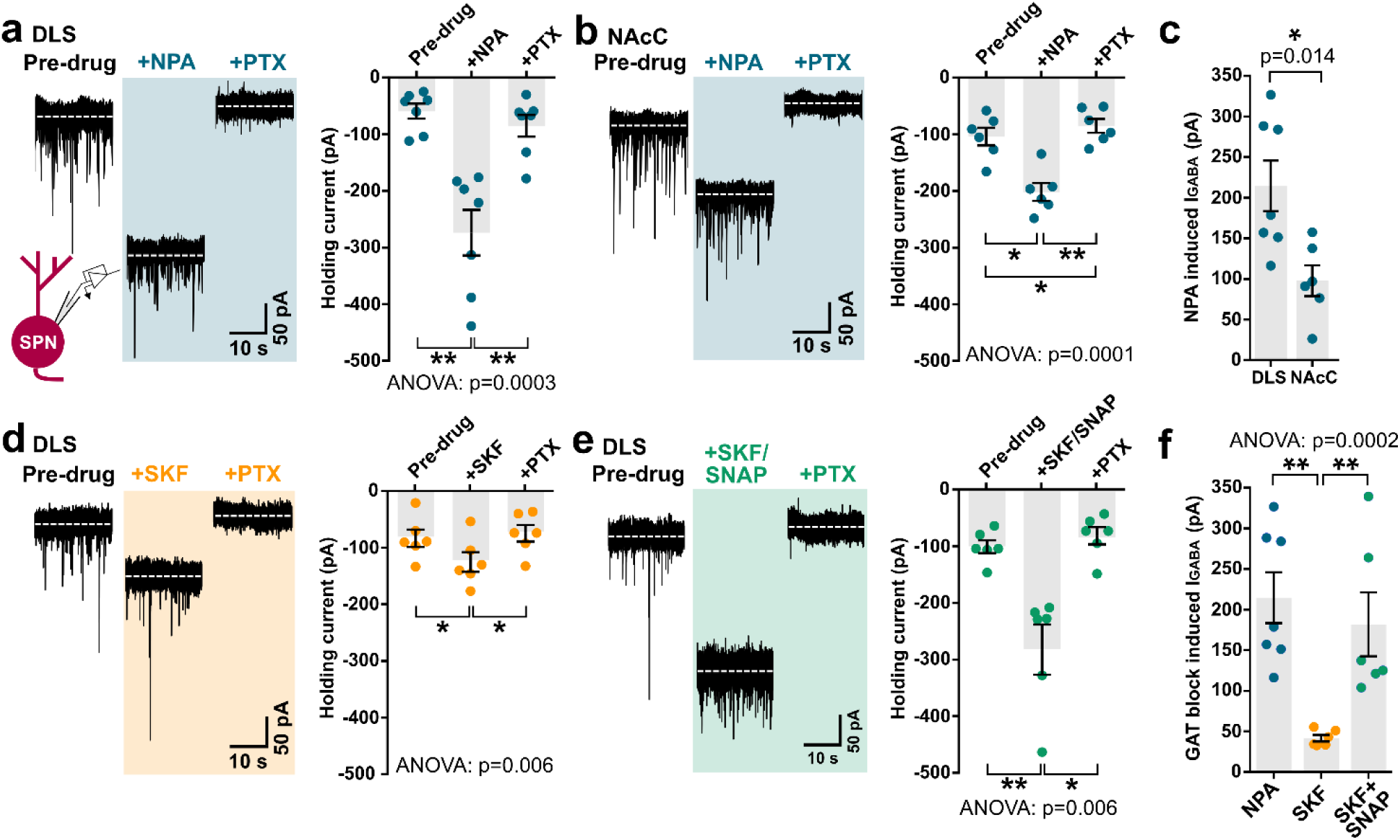
Tonic GABA currents in striatal spiny projection neurons (SPNs) are augmented by GAT inhibition. (**a**-**b, d**-**e**) *Left*, representative continuous whole-cell recordings from SPNs in DLS (a, d-e) or NAcC (b) voltage-clamped at −70 mV in the presence of ionotropic glutamate receptor antagonists NBQX (5 μM) and D-AP5 (50 μM), before and during bath application of (a-b) GAT inhibitor NPA (*blue*, 1.5 mM, n = 7 cells/5 mice for DLS in a, n = 6 cells/3 mice for NAcC in b), (d) GAT-1 specific inhibitor SKF89976A (*orange*, 20 μM, n = 6 cells/3 mice), or (e) the combined application of SKF89976A and GAT-3 specific inhibitor SNAP5114 (*green*, 50 μM, n = 6 cells/4 mice). GAT inhibitors increase the extracellular GABA_A_-mediated inward current, revealed by a shift in the holding current, and is reversed upon application of GABA_A_ receptor antagonist picrotoxin (PTX, 100 μM). *Right*, mean (± SEM) holding current in pA recorded in SPNs in control conditions, upon addition of GAT inhibitors and then PTX. (**c, f**) Mean (± SEM) tonic GABA_A_-receptor-mediated currents induced by GAT inhibition recorded from SPNs, calculated by subtracting pre-drug holding current from GAT block-induced holding current. Friedman’s ANOVA on Ranks and Student-Newman-Keuls multiple comparisons (a-b, d-e), Mann-Whitney U test (c), Kruskal-Wallis test and Dunn’s multiple comparisons (f). \**p* < 0.05, \*\**p* < 0.01.

We found also that tonic GABA inhibition of SPNs in DLS, like DA release, was regulated by both GAT-1 and GAT-3. Inhibition of GAT-1 alone with SKF89976A (20 μM) induced a small increase in the GABA_A_-mediated holding current (**Fig. 3d**; p = 0.006, Friedman’s ANOVA on Ranks, SKF vs. drug-free baseline: p = 0.001, SKF + PTX vs. drug-free baseline: p = 0.41, SKF vs. SKF + PTX: p = 0.01, Student-Newman-Keuls tests, n = 6 cells/3 mice). Combined inhibition of GAT-1 and GAT-3 with SKF89976A (20 μM) and SNAP5114 (50 μM) induced a three-fold increase (**Fig. 3e**; p = 0.006, Friedman’s ANOVA on Ranks, SKF + SNAP vs. drug-free baseline: p = 0.001, SKF + SNAP + PTX vs. drug-free baseline: p = 0.41, SKF + SNAP vs. SKF + SNAP + PTX: p = 0.011, Student-Newman-Keuls tests, n = 6 cells/4 mice), which was greater than after GAT-1 inhibition alone, but similar to that seen with broad-spectrum GAT inhibition NPA (**Fig. 3f**; p = 0.0002, Kruskal-Wallis ANOVA; SKF + SNAP vs. SKF: p < 0.01, NPA vs. SKF + SNAP: p > 0.05, NPA vs. SKF: p < 0.01; Dunn’s multiple comparison tests). These effects of GAT inhibition were due to GATs limiting an action potential-independent GABA tone i.e. due to “spontaneous” GABA release (Wójtowicz et al., 2013) since in the presence of Na_v_ blocker tetrodotoxin (TTX, 1 μM), NPA increased the GABA_A_-mediated holding current in SPNs in the DLS to a similar level to that induced in TTX-free conditions (**Supplementary Fig. 4**).

Collectively, these results show that striatal GAT-1 and GAT-3 regulate an ambient GABA tone, and to a greater degree in DLS than in NAcC. We explored an anatomical basis for this regional heterogeneity in GAT function. Striatal immunoreactivity to GAT-1 and GAT-3 in the DLS and NAcC revealed a modest relative enrichment in the DLS for both GAT-1 (**Fig. 4a-b**; p = 0.0093, Wilcoxon signed-rank test, n = 12 hemispheres/6 mice) and GAT-3 (**Fig. 4c-d**; p = 0.0015, Wilcoxon signed-rank test, n = 12 hemispheres/6 mice). We also noted enriched GAT-3 in the medial NAc shell (NAcS) contiguous with the medial septal nucleus (**Supplementary Fig. 5a-d**). This observation prompted us to test the effects of GAT inhibition on DA release in NAcS. Correspondingly, GAT inhibition diminished electrically evoked [DA]_o_ in NAcS to a greater degree than in NAcC (**Supplementary Fig. 5e-g**), indicating further regional heterogeneity in the role of GATs in limiting tonic inhibition.

**Fig 4.**
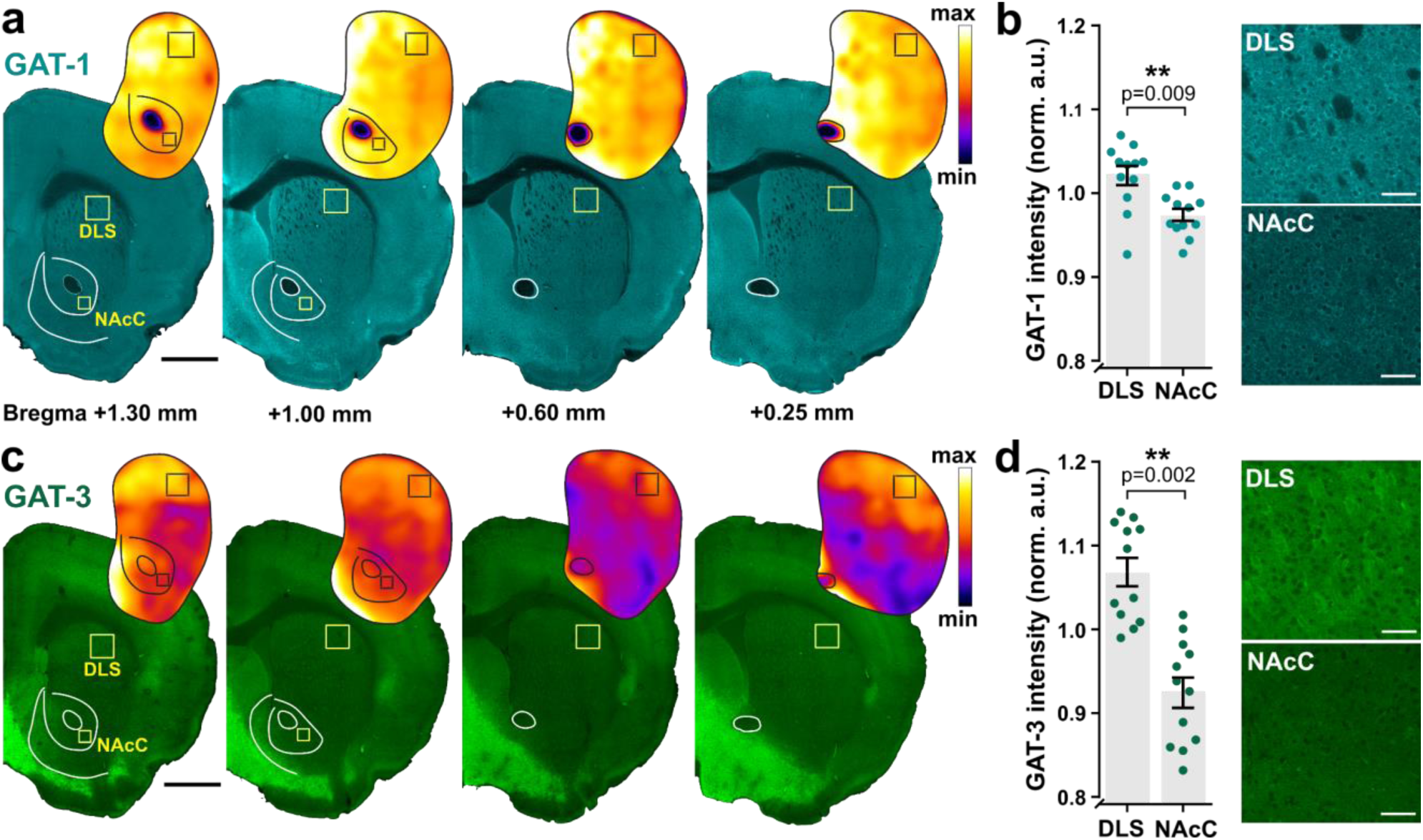
Enrichment of GAT-1 and GAT-3 expression in the DLS versus NAcC. (**a, c**) Representative immunofluorescence signals for GAT-1 (*cyan*, a) and GAT-3 (*green*, c) using confocal microscopy in coronal sections across the rostral-caudal limits containing striatum prepared from an individual C57BL/6J mouse with heat maps for striatal GAT intensity. Boxes indicate representative locations for GAT intensity measurements in the dorsolateral striatum (DLS) and nucleus accumbens core (NAcC). Scale bars: 1 mm. Note enriched GAT-3 in the medial NAc shell (NAcS) contiguous with the medial septal nucleus and enriched GAT-3 expression in the claustrum. (**b, d**) *Left*, Mean (± SEM) GAT-1 (b) and GAT-3 (d) intensity in DLS and NAcC normalized to total striatum and averaged across rostral-caudal sites for each hemisphere (n = 12 hemispheres/6 mice for each GAT-1 and GAT-3). *Right*, Representative single plane images of GAT-1 (b) and GAT-3 (d) immunofluorescence from DLS and NAcC; imaging parameters were kept constant across regions. Scale bars: 50 μm. Mann-Whitney U tests (b, d). \*\**p* < 0.01.

### GAT-1 and GAT-3 on astrocytes are key regulators of ambient GABA inhibition of DA release

Striatal GATs are located on the plasma membranes of cells that include GABAergic neurons (Augood et al., 1995; Durkin et al., 1995; Ng et al., 2000; Yasumi et al., 1997) and astrocytes (Chai et al., 2017; Ng et al., 2000; Yu et al., 2018). GATs have also been presumed, but not confirmed, to reside on DA axons to support GABA uptake, co-storage and co-release (Tritsch et al., 2014). To better understand where GATs are located to regulate tonic GABAergic inhibition of DA release, we probed two candidate locations, namely DA axons, and astrocytes. We explored whether GAT-1 or GAT-3 could be detected on DA axons using immunofluorescence and confocal microscopy, but we did not find robust evidence to support their localization on DA axons conditionally expressing an eYFP reporter (**Supplementary Fig. 6**). As a positive control for GAT-1 detection, we confirmed that GAT-1 could be localized to the neurites of parvalbumin (PV)-expressing GABAergic interneurons (**Supplementary Fig. 7**), which express GAT-1 (Augood et al., 1995).

In many brain regions, including striatum, astrocytes are thought to regulate ambient GABA levels through uptake (Yu et al., 2018). GAT-3 protein expression has been documented on striatal astrocytes (Chai et al., 2017; Ng et al., 2000; Yu et al., 2018), and although GAT-1 is typically associated with neuronal structures (Borden, 1996), recent transcriptomic studies have found RNA for both GAT-1 and GAT-3 in striatal astrocytes (Chai et al., 2017; Gokce et al., 2016; Zhang et al., 2014). We revisited GAT localization to astrocytes, using immunofluorescence and confocal microscopy with antibodies directed against either GAT-1 or GAT-3, as well as against the striatal astrocytic marker S100β (Chai et al., 2017) (**Fig. 5a-b**) in the DLS and NAcC. As expected, GAT-3 could be co-localized to S100β-expressing astrocytes (**Fig. 5d**), where GAT-3-immunoreactivity was observed distributed over a large surface area of plasma membrane when assessed in three dimensions (**Supplementary Fig. 8d-f**). We also found several instances of similar localization of GAT-1-immunoreactivity on S100β-expressing astrocytes (**Fig. 5c, Supplementary Fig. 8a-c**). These data indicate that GAT-3 and GAT-1 proteins can be expressed on the plasma membranes of striatal astrocytes.

**Fig 5.**
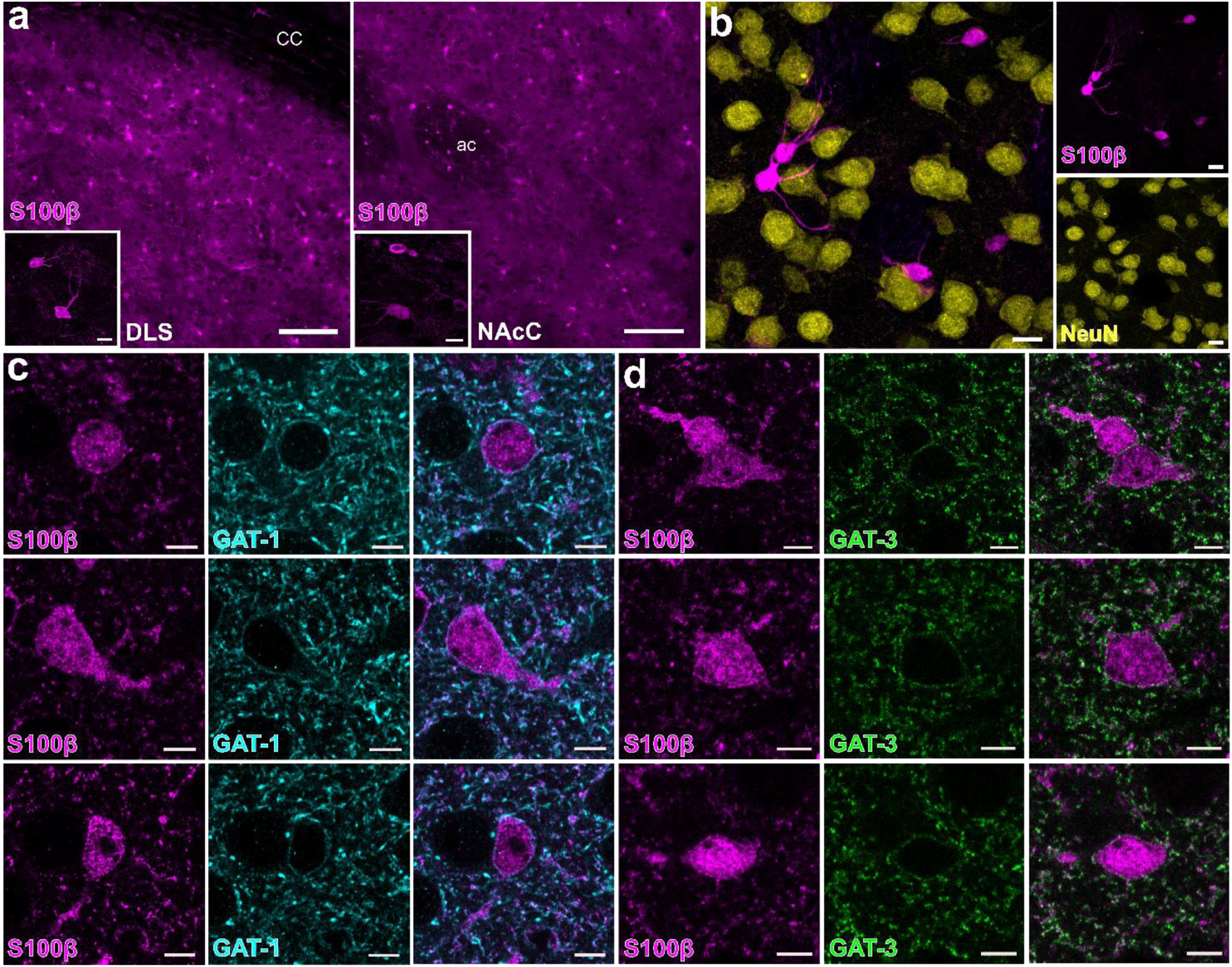
GAT-1 and GAT-3 are expressed on plasma membranes of striatal astrocytes. (**a**) Striatal immunofluorescence signals for astrocyte marker S100β (magenta) in dorsolateral striatum (DLS) and nucleus accumbens core (NAcC). Scale bars: 100 µm, for inset: 10 µm. *cc*: corpus callosum, *ac*: anterior commissure. (**b**) Immunofluorescence signals for S100β do not co-localize with immunofluorescence signals for neuronal marker NeuN. Scale bars: 10 µm. (**c**-**d**) GAT-3 (*green*, c) and GAT-1 (*cyan*, d) are expressed on plasma membranes of striatal S100β-expressing astrocytes imaged in DLS. Immunoreactivity for *left*, S100β, *centre*, GAT, *right*, merged. Scale bars: 5 μm.

Given the presence of GAT-3 and GAT-1 on striatal astrocytes, we probed whether astrocytes participate in regulating the level of inhibition of DA release by GABA. We exposed striatal slices to the gliotoxin fluorocitrate, which inhibits the enzyme aconitase, in turn disrupting the tricarboxylic acid cycle and inducing metabolic arrest in astrocytes (Henneberger et al., 2010; Martín et al., 2007; Paulsen et al., 1987). This approach has previously been established to render astrocytes inactive and prevent the effects of astrocytic GAT (Boddum et al., 2016; Bonansco et al., 2011). We pre-treated slices with fluorocitrate (200 µM for 45-60 min) or vehicle and then co-incubated with or without NPA (1.5 mM for 30 min), and assessed effect on [DA]_o_ evoked by 1p electrical stimulation across a range of sites in the DLS. We first confirmed that we could detect the effects of GAT inhibition in DLS in vehicle-treated control slices. Accordingly, [DA]_o_ evoked from slices incubated in NPA was significantly less than those incubated in NPA-free control conditions, as expected (**Fig. 6a**; U = 116, p = 0.00003, Mann-Whitney test, n = 24 observations/5 mice for each condition), and the 4p/1p ratio (50 Hz) was appropriately enhanced (**Fig. 6b**; t_(14)_ = 2.988, p = 0.009; unpaired Student’s t-test, n = 8 experiments/4 mice for each condition). By contrast, when we performed these experiments in slices pre-treated with fluorocitrate to inactivate astrocytes, NPA did not significantly modify [DA]_o_ evoked by 1p (**Fig. 6c**; U = 699, p = 0.103, Mann-Whitney test, n = 42 observations/7 mice for each condition), or the 4p/1p ratio (50 Hz), compared to NPA-free conditions (**Fig. 6d**; t_(24)_ = 0.5384, p = 0.595; unpaired Student’s t-test, n = 13 experiments/7 mice for each condition). The effect of NPA on [DA]_o_ when astrocytes were inhibited was significantly less than when astrocytes were intact (**Fig. 6e;** U = 288, p = 0.0036, Mann-Whitney test). We also noted that a comparison of evoked [DA]_o_ with and without fluorocitrate revealed that [DA]_o_ were reduced by fluorocitrate treatment (**Fig. 6f**; U = 226, p = 0.0001, Mann-Whitney test). Together, these data suggest that astrocytic GATs support GABA uptake and limit the level of inhibition of DA release by ambient GABA, such that in turn, astrocytes indirectly support DA release.

**Fig 6.**
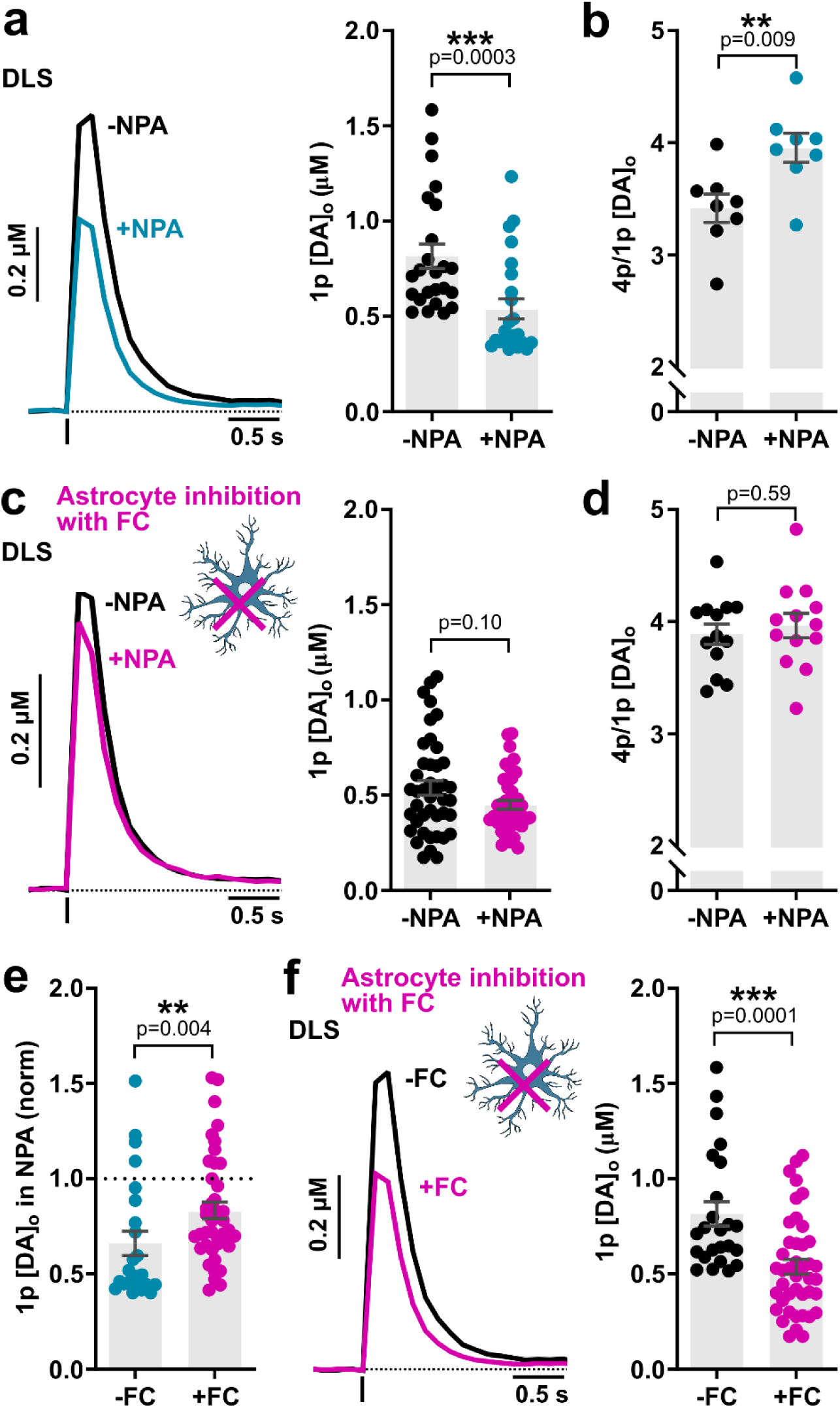
GATs on striatal astrocytes regulate GABA inhibition of DA release. (**a**-**d**) Mean profiles of [DA]_o_ and mean peak [DA]_o_ (± SEM) in DLS evoked by 1 electrical pulse (a, c) or 4 pulses normalized to 1p (b, d) in the absence (*black*) and presence of GAT inhibitor NPA (*blue* or *pink*, 1.5 mM) in vehicle-treated control slices (a-b, n = 5 mice) or in slices treated with astrocyte inhibitor fluorocitrate (FC, 200 µM) (c-d, n = 7 mice). (**e**) Mean peak [DA]_o_ (± SEM) evoked by 1 electrical pulse in the presence of NPA (1.5 mM) normalized to control conditions in control slices (*blue*) or fluorocitrate-treated slices (*pink*). (**f**) Mean peak [DA]_o_ (± SEM) evoked by 1 electrical pulse in the absence of NPA in control slices (*-FC*) and in fluorocitrate-treated slices (+FC) from (a, b). DHβE (1 μM) present throughout. Mann-Whitney U tests (a, c, e, f) and Student’s unpaired t-tests (b, d). \*\**p* < 0.01, \*\*\**p* < 0.001

### Tonic inhibition of DA release in the DLS is augmented in a mouse model of parkinsonism

Our data provide compelling evidence that GAT function regulates DA output level in the DLS. Dysregulation of GATs in the basal ganglia has now been implicated in several models of neurological disease: in 6-OHDA toxin-based rat and mouse models of dopamine depletion in Parkinson’s, astrocytes in the external globus pallidus have downregulated GAT-3 (Chazalon et al., 2018); and in R6/2 and FVB/N transgenic mouse models of Huntington’s disease, striatal GAT expression is increased and tonic inhibition by ambient GABA decreased (Cepeda et al., 2013; Wójtowicz et al., 2013; Yu et al., 2018). Given that deficits in DA transmission in DLS, but not in NAcC, are common to transgenic rodent models of parkinsonism prior to cell loss (Janezic et al., 2013; Sloan et al., 2016; Taylor et al., 2014), we explored whether tonic GABAergic inhibition of striatal DA release and its regulation by striatal GATs might be dysregulated in a mouse model of early parkinsonism.

We chose the *SNCA*-OVX mice, a BAC transgenic mouse model of early parkinsonism (Janezic et al., 2013). *SNCA*-OVX mice are devoid of mouse α-synuclein but overexpress human wildtype α-synuclein at disease-relevant levels and show early deficits in DA release prior to DA cell loss (Janezic et al., 2013). To address our aims, we made these *SNCA*-OVX mice ‘optogenetics capable’, such that they allowed for optical manipulation of DA axons, and we first characterized neurotransmission from DA axons in these mice. We crossed *Slc6a3*^*IRES-Cre*^ mice with α-synuclein knockout mice to create *Slc6a3*^*IRES-Cre*^ mice devoid of mouse α-synuclein, then crossed these mice with S*NCA*-OVX mice to generate two genotypes of mice, both devoid of mouse α-synuclein: (1) “*SNCA*+” mice that express Cre recombinase in DA neurons and human α-synuclein; and (2) as littermate controls, “*Snca-/-*” mice, that express Cre recombinase in DA neurons but no human transgene. We confirmed that, as observed in the original *SNCA*-OVX mice (Janezic et al., 2013), the resulting *SNCA*+ mice at 4 months of age exhibited a ∼30% deficit in electrically evoked [DA]_o_ when compared to littermate controls (*Snca*-/-) in the dorsal striatum (**Fig. 7a**; t_(46)_ = 3.272, p = 0.0020; unpaired Student’s t-test, n = 24 observations/5 mice for each genotype) but not in the NAc (**Fig. 7a**; t_(40)_ = 1.393, p = 0.1714; unpaired Student’s t-test; n =21 observations/5 mice for each genotype). The DA release deficit was also not attributable to a reduction in striatal DA content in *SNCA*+ mice compared to *Snca*-/- mice (**Fig. 7b**; dorsal striatum: t_(14)_ = 0.1625, p = 0.8733, n = 8 experiments/5 mice for each genotype; NAc: t_(14)_ = 0.7445, p = 0.4689, unpaired Student’s t-tests; n = 8 experiments/5 mice for each genotype) establishing an underlying change to DA releasability rather than storage potential. We then verified that [DA]_o_ evoked optogenetically in DLS by single light pulses showed a similar deficit in *SNCA*+ compared to *Snca-/-* (**Fig. 7c-d**; t_(29)_ = 2.443, p = 0.0209, unpaired Student’s t-test; F_(1,12)_ = 7.108, p = 0.0206; two-way repeated-measures ANOVA; *SNCA+*: n = 16 observations/3 mice; *Snca-/-*: n = 15 observations/3 mice). Having established a deficit in DA release in this optogenetics-capable mouse model of parkinsonism, we next addressed for the first time whether DA release deficits are accompanied by corresponding deficits in GABA co-release from DA axons. Using voltage-clamp recordings in SPNs, we observed a significantly lower amplitude of IPSCs evoked by light-activation of DA axons in *SNCA+* compared to *Snca*-/- mice (**Fig. 7e-f**; t_(14)_ = 2.680, p = 0.0179, unpaired Student’s t-test; F_(1,14)_ = 7.281, p = 0.0173; two-way repeated-measures ANOVA; *SNCA+*: n = 7 cells/4 mice; *Snca-/-*: n = 9 cells/4 mice) indicating a companion deficit in GABA release, and these IPSCs exhibited a similar gradual run-down to DA release (**Fig. 7d**,**f**). Light-evoked IPSCs were GABA_A_ receptor-mediated as they were eliminated by picrotoxin (PTX, 100 µM) (**Supplementary Fig. 9a**,**b**) and the observed differences in IPSC amplitudes were not due to differences in series resistance (**Supplementary Fig. 9c**).

**Fig 7.**
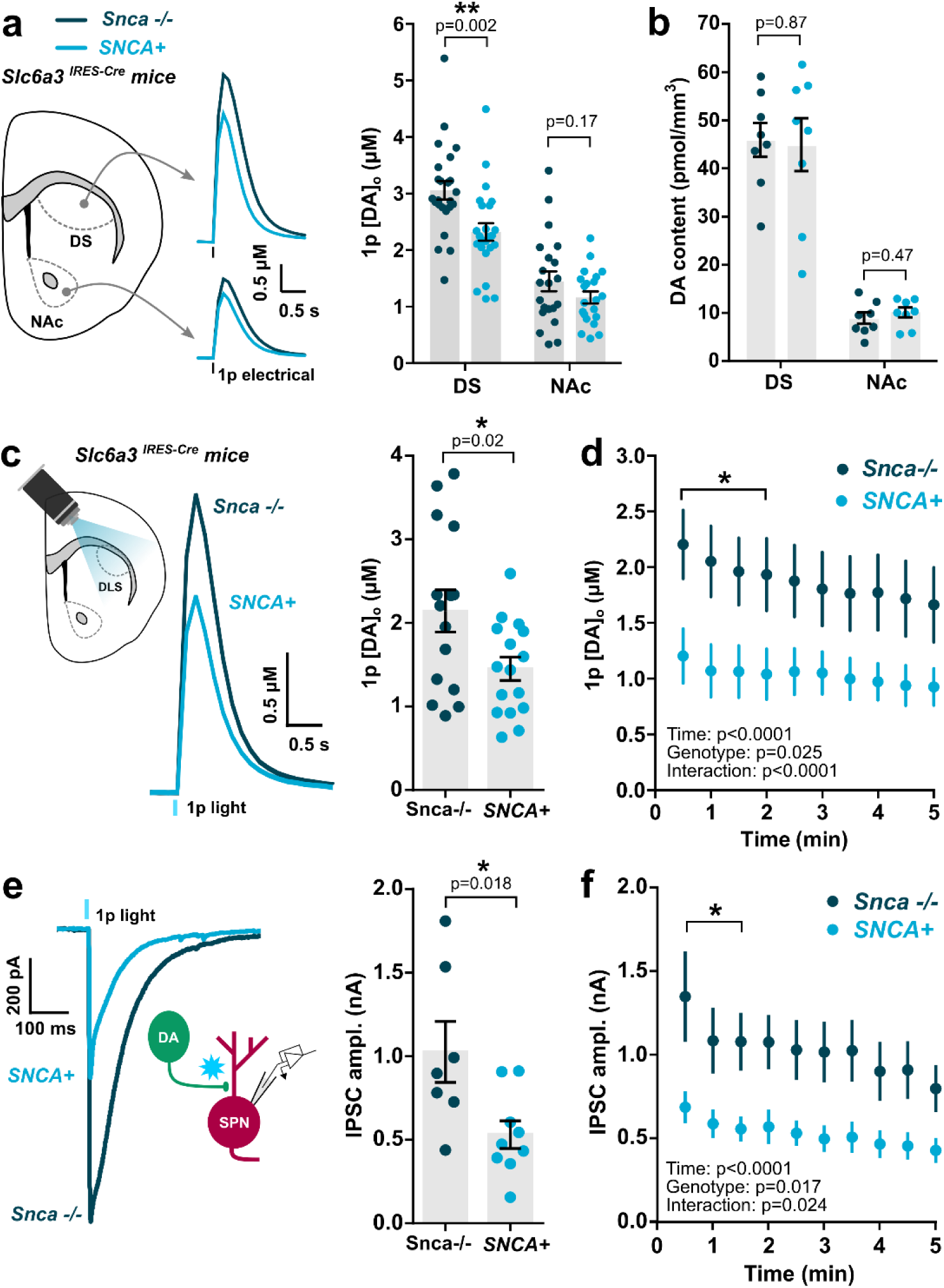
Attenuated GABA and DA co-release from DA axons in a transgenic mouse model of early parkinsonism. (**a**) *Left*, mean [DA]_o_ profiles vs. time following 1-pulse electrical simulation in dorsal striatum (DS) and nucleus accumbens (NAc) of *SNCA*+ mice (*light blue*) and littermate controls (*Snca*-/-, *dark blue*) at 3-4 months, backcrossed onto an *Slc6a3*^*IRES-Cre*^ background. *Right*, Mean (± SEM) 1p-evoked [DA]_o_ (in μM) from *Left* (n = 24 observations/5 mice per genotype in DS, n = 21 observations/5 mice per genotype in NAc). (**b**) Mean (± SEM) DA content in DS and NAc of *SNCA*+ mice (*light blue*) and littermate controls (*Snca*-/-, *dark blue*) (n = 8 experiments/5 mice per genotype in DS and NAc). (**c**) *Left*, mean [DA]_o_ profiles vs. time following 1-pulse light simulation in DLS of *SNCA*+ mice (*light blue*) and littermate controls (*Snca*-/-, *dark blue*). *Right*, Mean (± SEM) and individual values for 1p-evoked [DA]_o_ (in μM) from *Left* (n = 15 observations/3 mice in *Snca*-/- mice, n = 16 observations/3 mice in *SNCA*+ mice). (**d**) Mean (± SEM) 1p light-evoked [DA]_o_ (in μM) recorded every 30 s in DLS of *SNCA*+ mice (*light blue*, n = 7 experiments/3 mice) and littermate controls (*Snca*-/-, *dark blue*, n = 7 experiments/3 mice). (**e**-**f**) Example, mean (± SEM) and individual values for 1-pulse light-evoked inhibitory postsynaptic currents (IPSCs) recorded from spiny projection neurons (SPNs) every 30 s in the DLS of *SNCA+* mice (*light blue*, n = 9 cells/4 mice) and littermate controls (*Snca*-/-, *dark blue*, n = 7 cells/4 mice), voltage clamped at −70 mV and in the presence of ionotropic glutamate receptor antagonists (NBQX, 5 μM; D-APV, 50 μM). Student’s unpaired t tests (a-c, e) and two-way repeated-measures ANOVA with Sidak’s multiple comparison tests (d, f). \**p* < 0.05, \*\**p* < 0.01.

We then explored in this model whether tonic GABA inhibition of DA release was modified in DLS or NAcC. We found that GABA_R_ antagonism enhanced [DA]_o_ evoked by single light pulses to a significantly greater degree in *SNCA+* mice than in *Snca-/-* controls in DLS (**Fig. 8a**; F_(1,13)_ = 12.42, p = 0.0037; two-way repeated-measures ANOVA; *SNCA+*: n = 8 experiments/5 mice; *Snca-/-*: n = 7 experiments/5 mice) but not in NAcC (**Fig. 8b**; F_(1,15)_ = 2.318, p = 0.1487; two-way repeated-measures ANOVA; *SNCA+*: n = 8 experiments/5 mice; *Snca-/-*: n = 9 experiments/5 mice), which was a significant regional difference (**Fig. 8c-d**; p < 0.0001, Kolmogorov-Smirnov test; U = 10, p = 0.0207, Mann-Whitney test). These data indicate that tonic GABA inhibition of DA axons is elevated in the DLS of *SNCA+* mice.

**Fig 8.**
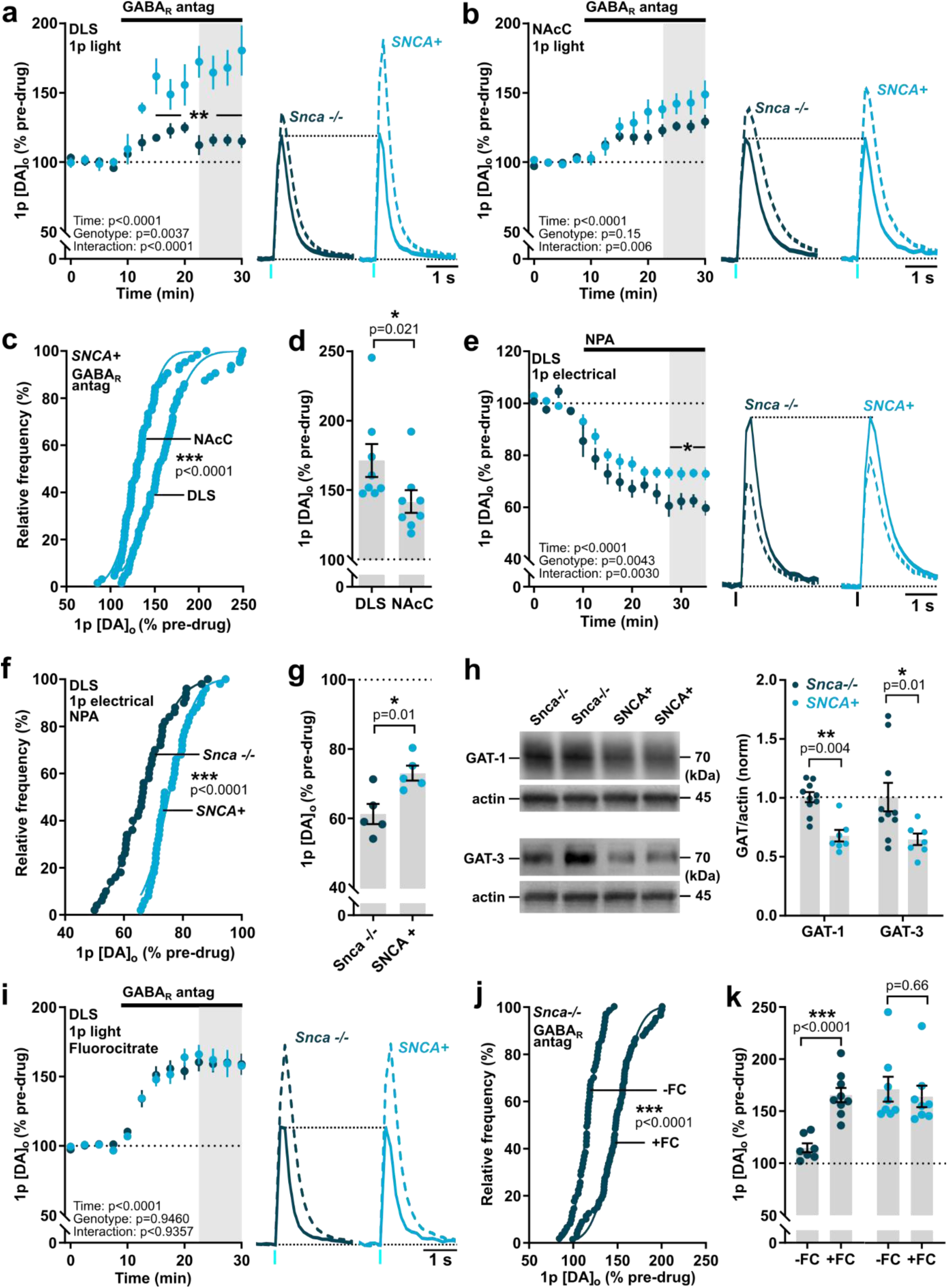
Enhanced tonic inhibition of striatal DA release and impaired GAT function in mouse model of early parkinsonism. (**a**,**b**,**e**,**i**) Mean peak [DA]_o_ (± SEM) during consecutive recordings evoked by 1p light (a,b,i) or electrical pulse (e) in DLS (a,e,i) or NAcC (b) during applications of antagonists for GABA_A_ (bicuculline, 10 µM) and GABA_B_ receptors (CGP 55845, 4 µM) (a,b,i), or the nonspecific GAT inhibitor NPA (1.5 mM)(e) in slices pre-incubated (i) or not pre-incubated (a,b,e) with fluorocitrate (FC, 200 µM, 45-60 min) from *Snca*-/- (*dark blue*, GABA_R_ antagonism: n = 7 experiments/5 mice in DLS (a), n = 9 experiments/5 mice in NAcC (b), n = 7 experiments/3 mice for fluorocitrate (i); NPA: n = 5 experiments/4 mice (e)) and *SNCA*+ mice (*light blue*, GABA_R_ antagonism: n = 8 experiments/5 mice for both DLS and NAcC (a,b), n = 6 experiments/3 mice for fluorocitrate (i); NPA: n = 5 experiments/4 mice (e)). Data are normalized to mean of 4 time points prior to drug application (*dotted line*); mean transients of [DA]_o_ are derived from last 4 time points (*gray shaded region*) and normalized to pre-drug baselines. (**c-d**,**f-g**,**j-k**) Cumulative frequency plots of individual data points (c,f,j) and mean (± SEM) per recording site *(d,g,k*) from (a,b,e,i). (**h**) Representative Western blots and mean (± SEM) of GAT-1 and GAT-3 protein content of dorsal striatum tissue taken from *Snca*-/- mice (n = 10 mice) and *SNCA*+ mice (n = 7 mice). Data normalized to actin and littermate control expression. Two-way repeated-measures ANOVA with Sidak’s multiple comparison tests (a,b,e,i), Komogorov-Smirnov tests (c,f,j), Student’s unpaired t-tests (d,g,k) and Mann-Whitney U tests (h). \**p* < 0.05, \*\**p* < 0.01, \*\*\**p* < 0.0001.

We tested the hypothesis that elevated tonic inhibition of DA release in the DLS of *SNCA+* mice might be due to impaired GAT function. We identified that there was an impairment in the effect of the non-selective GAT inhibitor NPA on DA release in DLS: there was an attenuated effect of NPA on [DA]_o_ evoked by single electrical pulses in *SNCA+* versus *Snca- /-* controls (**Fig. 8e-g**; F_(1,8)_ = 5.790, p = 0.0428; two-way repeated-measures ANOVA; p < 0.0001, Kolmogorov-Smirnov test; t_(8)_ = 3.244, p = 0.0118, unpaired Student’s t-test; *SNCA+*: n = 5 experiments/4 mice; *Snca-/-*: n = 5 experiments/4 mice). Furthermore, quantification of Western blots of dorsal striatal tissue revealed significantly lower levels of both GAT-1 and GAT-3 proteins in *SNCA+* mice versus *Snca-/-* controls (**Fig. 8h**; GAT-1: U = 2, p = 0.0004; GAT-3: U = 10, p = 0.0136, Mann-Whitney tests; n = 7 *SNCA+* mice, n = 10 *Snca-/-* mice). Given the expression and role of both GATs on astrocytes in regulating DA release, these data strongly implicate GAT downregulation on astrocytes as a contributing cause of enhanced GABA inhibition of DA release in *SNCA*+ mice. We therefore tested whether by impairing astrocytic GAT function through astrocyte inactivation with fluorocitrate, we might equalise the level of tonic GABA inhibition operating in the DLS of *SNCA*+ and *Snca*-/- mice. In slices pre-incubated with fluorocitrate (200 µM for 45-60 min), the difference in the effect of GABA_R_ antagonists on [DA]_o_ evoked by single light pulses seen in DLS between *SNCA+* mice and *Snca-/-* controls was completely abolished, and the effect of GABA_R_-antagonists was not different between genotypes (**Fig. 8i**; F_(1,15)_ = 0.005, p = 0.946; two-way repeated-measures ANOVA; *SNCA+*: n = 8 experiments/3 mice; *Snca-/-*: n = 9 experiments/3 mice). Fluorocitrate incubation significantly boosted the effect of GABA_R_ antagonists in *Snca-/-* mice (**Fig. 8j-k**; p < 0.0001, Kolmogorov-Smirnov test; t_(14)_ = 5.82, p < 0.0001, unpaired Student’s t-test) but had no effect in *SNCA+* mice (**Fig. 8k**; t_(14)_ = 0.44, p = 0.664, unpaired Student’s t-test). Taken together, these data suggest that tonic inhibition of DA release by ambient GABA is augmented in the dorsal striatum in early parkinsonism due to downregulation of GATs that are located at least in part to astrocytes (**Fig. 9**).

**Fig 9.**
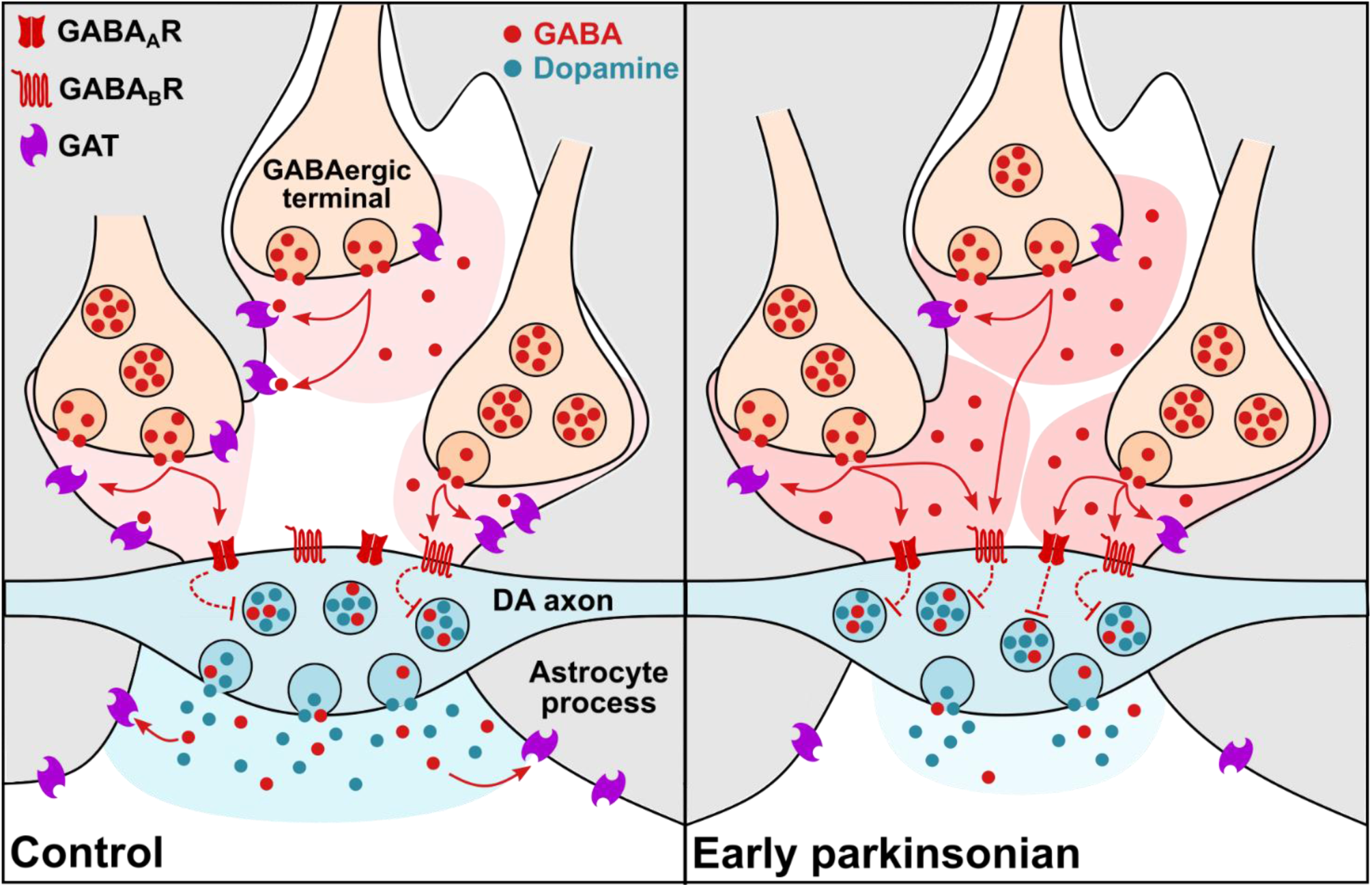
Augmented tonic inhibition of striatal DA release in dorsal striatum in early parkinsonism due to reduced striatal GAT expression. Under normal circumstances (*left*), GAD-synthesized GABA is released from GABAergic striatal neurons can spillover to act at GABA receptors (GABA_A_R and GABA_B_R) located presumably on DA axons, inhibiting (*dashed red lines*) DA and GABA co-release. The level of GABA spillover and tonic inhibition of DA release is determined by the activity of GABA transporters (GATs) located on astrocytes (*gray*) and neurons, which remove GABA from the extracellular space. In a mouse model of early Parkinsonism (*right*), striatal GAT expression is downregulated in dorsal striatum, resulting in augmented tonic inhibition of DA release by GABA. Co-release of GABA from DA axons is also reduced.

## DISCUSSION

We define a major role for striatal GATs and astrocytes in setting the level of DA output in the striatum. We show that GAT-1 and GAT-3, located at least in part on striatal astrocytes, govern tonic GABAergic inhibition of DA release. GATs operate in a heterogeneous manner across the striatum, substantially limiting tonic inhibition of DA release in DLS but not NAcC. Moreover, in a mouse model of early parkinsonism, we reveal maladaptive decreases in striatal GAT-1 and GAT-3 expression and consequently, profound augmentation of tonic inhibition of DA release by GABA in the dorsal striatum.

### GATs limits the tonic inhibition of DA release

We found that tonic inhibition of DA release by GABA spans dorsal-ventral territories of striatum, and arises from a GAD-dependent source of GABA. The source of GABA was not a non-canonical ALDH-dependent source e.g. co-release from DA axons, as inhibition of ALDH did not attenuate the tonic inhibition of DA release by GABA, despite attenuating GABA co-release from DA axons, as seen previously (Kim et al., 2015). Conversely, ALDH-inhibition even slightly boosted tonic inhibition of DA release, suggesting that ALDH-dependent sources of GABA, such as GABA co-release from DA axons, limit the tonic inhibition by the GAD-dependent GABA network. Correspondingly, in mice overexpressing human α-synuclein, in which we found that GABA co-release from DA axons is attenuated, we also found that the levels of tonic GABA inhibition on DA release was boosted. We note that *Aldh1a1* mutations in humans and deletion in mice lead to alcohol-consuming preferences (Kim et al., 2015; Liu et al., 2011; Sherva et al., 2009), and speculate that dysregulated DA output might plausibly result and contribute to this behaviour.

The paucity of GABAergic synapses on DA axons (Charara et al., 1999) suggests that GAD-dependent GABA tone arises from the extrasynaptic ambient tone that can be detected in striatum (Ade et al., 2008; Cepeda et al., 2013; Kirmse et al., 2008, 2009; Santhakumar et al., 2010). This tone was action potential independent, i.e. spontaneous (Kaeser and Regehr, 2013), as reported previously for tonic inhibition of SPNs (Wójtowicz et al., 2013). A spontaneous GABAergic regulation of DA release is not surprising when considering that the axonal arbour of a given nigrostriatal DA neuron (in rat) reaches on average 2.7% of the volume of striatum (Matsuda et al., 2009; Oorschot, 1996), and that such volumes contain ∼74,000 GABAergic neurons (calculated from 2.8 million striatal neurons per hemisphere (Oorschot, 1996), of which ∼98% are GAD-immunoreactive) and also GAD-positive cholinergic interneurons that can corelease GABA (Lozovaya et al., 2018). Even very low rates of spontaneous vesicle release from a small fraction of GAD-utilizing GABAergic neurons might summate sufficiently to provide a tone at GABA receptors on DA axons that limits DA output. The general functions of this spontaneous GABA tone are not well understood, but could differ from functions of action potential-dependent or synaptic events (Farrant and Nusser, 2005), and could include regulation of DA axonal membrane resistance to modify the impact of other inputs or limit the propagation of action potentials through the axonal arbour for a sparser coding.

We found that GAT-1 and GAT-3 both limit the actions of GABA on DA axons in DLS, and thereby indirectly facilitate DA release. This unprecedented role for the GATs in supporting DA output was heterogeneous: GATs limited tonic GABAergic inhibition of DA release in DLS, and markedly less so in NAcC, which corresponded with heterogeneity in GAT-1 and GAT-3 expression. Of note, the positive relationship we find between GAT function and DA output is paralleled by, and provides a candidate explanation for, some clinical effects of GAT inhibitors e.g. tiagibine. When used clinically as anti-epileptics to increase extracellular GABA levels, these inhibitors can have parkinsonian-like motor side effects (Zaccara et al., 2004).

We did not find evidence for robust localization of GAT-1 or GAT-3 proteins to DA axons in DLS, despite a previous inference that GATs reside on DA axons to support GABA uptake for co-release (Tritsch et al., 2014). This inference was based on detection of mRNA for GAT-1 (and weakly for GAT-3) in the somata of DA neurons in substantia nigra, and on the attenuation of GABA co-release from DA axons after pharmacological inhibition of both GAT-1 and GAT-3 (Tritsch et al., 2014). However, because subsequent work has shown that there is a tonic GABAergic inhibition of DA release mediated by both GABA_A_ and GABA_B_ receptors (Lopes et al., 2019), which we show here is profoundly limited by the GATs, then the observed dependence of GABA co-release on GAT could be explained alternatively by GATs limiting tonic inhibition of GABA co-release, rather than GATs necessarily being required for GABA uptake.

We revealed that astrocytes play a critical role in limiting the tonic inhibition of DA release and therefore supporting DA output. We found that both GAT-3 and, to a lesser extent, GAT-1, could be identified on astrocytes, challenging the long-held generalization that GAT-1 is exclusively neuronal (Borden, 1996). Furthermore, we found that astrocyte inactivation with the glial metabolic poison fluorocitrate prevented the effects of GAT inhibitors and boosted tonic GABAergic inhibition of DA release. Although the mechanisms and specificity of fluorocitrate actions are incompletely understood, and prolonged treatment with fluorocitrate has the potential to compromise neuronal integrity, this pharmacological approach remains one of the few tools available to render astrocytes and their transporters inactive. Furthermore, exposure over short durations (1 hr), as used here, is thought to have limited effects on downstream neuron viability (Henneberger et al., 2010; Martín et al., 2007). The role we find for astrocytes in supporting GABA uptake to limit tonic inhibition of DA release, indicates a previously unappreciated role for astrocytes in determining the dynamics of DA signaling. This finding significantly revises current understanding of the striatal mechanisms that can dynamically regulate DA transmission. Astrocytic GATs have recently been shown to regulate tonic GABAergic inhibition of striatal SPNs and striatal-dependent behaviors (Yu et al., 2018), and thus, our collective findings point to GATs and astrocytes as powerful regulators of striatal and DA function that warrant further future investigation.

### Striatal GAT dysfunction in a mouse model of Parkinson’s disease

To probe the wider potential significance of the regulation of striatal DA by striatal GATs, we explored GAT function in a mouse model of early parkinsonism. A recent study in external globus pallidus of dopamine-depleted rodents found elevated extracellular GABA resulting from downregulation of GAT-3 on astrocytes, mediated through a loss of DA signalling at D_2_ DA receptors (Chazalon et al., 2018). Conversely, striatal GAT-3 levels are upregulated in mouse models of Huntington’s disease (Wójtowicz et al., 2013; Yu et al., 2018). In an intriguing parallel seen for glutamate transmission in pre-neurodegenerative β-amyloid-based mouse models of early Alzheimer’s disease, hippocampal neurons become hyperactive due to an attenuation of glutamate uptake by astrocytes (Zott et al., 2019). Together these emerging strands suggest that impaired astrocyte transporters might be an early underlying feature across neurodegenerative diseases. We explored potential adaptations to GAT function and tonic GABA inhibition of DA release in the striatum of the human α-synuclein-overexpressing mouse model of PD. This model is a highly physiological, slowly progressing mouse model of parkinsonism, that, in capturing a human disease-relevant genetic burden of α-synuclein overexpression, shows early deficits in DA release restricted to dorsal striatum prior to degeneration of DA neurons, disturbed encoding of behaviour of surviving DA neurons and a motor phenotype in old age (Dodson et al., 2016; Janezic et al., 2013). We firstly ascertained the novel finding that DA transmission deficits in the DLS of this model in early adulthood are accompanied by a corresponding deficit in GABA co-release from DA axons. Furthermore, we found an augmentation of tonic GABA inhibition of DA release in the DLS (and not NAcC), which was accompanied by downregulated GAT-1 and GAT-3 expression. Whether these adaptations in GAT are consequential to reduced dopamine signalling, as occurs in astrocytes in globus pallidus after depletion of dopamine (Chazalon et al., 2018), or to reduced GABA co-release, or to a potential interaction between α-synuclein and striatal GATs and/or astrocytes is not yet known. Regardless, this resulting enhanced tonic inhibition will diminish nigrostriatal DA release, compounding the release deficits underpinned by α-synuclein e.g. tighter vesicle clustering at DA release sites (Janezic et al., 2013). These changes in GATs and tonic GABA inhibition in early parkinsonism can therefore be considered ‘maladaptive’ to DA loss.

In conclusion, the regulation of striatal GABA-DA interactions via striatal GATs and astrocytes represent loci for governing DA output as well as for maladaptive plasticity in early parkinsonism, which could also provide a novel therapeutic avenue for upregulating DA signalling in PD.

## MATERIALS AND METHODS

### Mice

All experimental procedures involving the use of animals were carried out according to institutional guidelines and conformed to the UK Animals (Scientific Procedures) Act 1986. Wild-type C57BL/6J mice were obtained from Charles River (Harlow, UK). Knock-in mice bearing an internal ribosome entry site (IRES)-linked Cre recombinase gene downstream of the gene *Slc6a3*, which encodes the plasma membrane dopamine transporter (DAT) were obtained from Jackson Laboratories (*Slc6a3*^*IRES-Cre*^ mice; *B6.SJL-Slc6a3*^tm1.1(cre)Bkmn^/J; stock no. 006660). PV^*Cre*^ knock-in mice expressing Cre recombinase in parvalbumin (PV)-expressing neurons were obtained from Jackson Laboratories (*B6;129P2-Pvalb*^tm1(cre)Arbr^/J; stock no. 008069). BAC-transgenic mice that overexpress human α-synuclein (*SNCA*) at Parkinson’s disease-relevant levels and are back-crossed onto an α-synuclein-null (*Snca*^−/−^) background (B6.Cg-Tg(*SNCA*)OVX37Rwm Snca^tm1Rosl^/J; Jackson Laboratories stock no. 023837), “*SNCA*-OVX” mice, were bred locally (Janezic et al., 2013). ‘Optogenetic capable’ *SNCA+* mice were generated by crossing *Slc6a3*^IRES-Cre^+/+; *Snca*-/- mice with *SNCA*+/-; *Snca*-/- mice (*SNCA*-OVX mice) (Janezic et al., 2013). For all experiments involving *SNCA*+ mice, we used age- and sex-matched *Snca*-null mice (heterozygous for Slc6a3^*IRES-Cre*^) as littermate controls. All mice were maintained on a C57BL/6 background, group-housed and maintained on a 12-hr light cycle with *ad libitum* access to food and water. All transgenic mice used in experiments were homozygous for transgenes or mutant alleles.

### Stereotaxic intracranial injections

Slc6a3^*IRES-Cre*^ mice, Slc6a3^IRES-Cre^x*SNCA*-OVX mice (postnatal day (P) 77-84) or PV^*Cre*^ mice (P28-35) were anesthetized with isoflurane and placed in a small animal stereotaxic frame (David Kopf Instruments). After exposing the skull under aseptic techniques, a small burr hole was drilled and adeno-associated virus (8×10^12^ genome copies per ml; UNC Vector Core Facility) encoding Cre-dependent ChR2 was injected. Viral solutions were injected at an infusion rate of 100 nL/min with a 32-gauge Hamilton syringe (Hamilton Company) and withdrawn 5-10 min after the end of injection. In Slc6a3^IRES-Cre^ x *SNCA*-OVX mice, and Slc6a3^IRES-Cre^ mice, a total volume of 1 μL of AAV5-EF1α-DIO-hChR2(H134R)-eYFP was injected bilaterally (500 nL per hemisphere/injection) into substantia nigra pars compacta (SNc, AP −3.1 mm, ML ±1.2 mm from bregma, DV −4.25 mm from exposed dura mater). In PV^*Cre*^ mice, a total volume of 600 nL of AAV2-EF1α-DIO-hChR2(H134R)-eYFP was injected bilaterally (300 nL per hemisphere/injection) into dorsolateral striatum (DLS, AP +0.65 mm, ML ±2.0 mm from bregma, DV −1.85 mm from exposed dura mater). Viral-injected mice were used for experiments after >28 days post-injection.

### Slice preparation

Acute brain slices were obtained from 35- to 80-day-old mice using standard techniques. Mice were culled by cervical dislocation (for FSCV experiments alone) or mice were anaesthetized with pentobarbital and transcardially perfused with ice-cold artificial cerebrospinal fluid (aCSF) containing (in mM): 130 NaCl, 2.5 KCl, 26 NaHCO_3_, 2.5 CaCl_2_, 2 MgCl_2_, 1.25 NaH_2_PO_4_ and 10 glucose (for whole-cell patch-clamp electrophysiology experiments alone or in combination with FSCV experiments). 300 μm-thick coronal slices containing caudate putamen (CPu) and NAc were prepared from dissected brain tissue using a vibratome (VT1200S, Leica Microsystems) and transferred to a holding chamber containing a HEPES-based buffer solution maintained at room temperature (20-22°C) containing (in mM): 120 NaCl, 20 NaHCO_3_, 10 glucose, 6.7 HEPES acid, 5 KCl, 3.3 HEPES sodium salt, 2 CaCl_2_, 2 MgSO_4_, 1.2 KH_2_PO_4_ (for FSCV experiments alone) or containing aCSF kept at 34°C for 15 min before returning to room temperature (20-22°C). All recordings were obtained within 5-6 hours of slicing. All solutions were saturated with 95% O_2_ / 5% CO_2_.

### Fast-scan cyclic voltammetry (FSCV)

Individual slices were hemisected and transferred to a recording chamber and superfused at ∼3.0 mL/min with aCSF at 31-33 °C. A carbon fibre microelectrode (CFM; diameter 7-10 μm, tip length 70-120 μm), fabricated in-house, was inserted 100 μm into the tissue and slices were left to equilibrate and the CFM to charge for 30-60 min prior to recordings. All experiments were carried out either in the dorsolateral quarter of the CPu of the striatum (DLS) or nucleus accumbens (NAc) core (NAcC; within 100 μm of the anterior commissure) or lateral NAc shell (NAcS), one site per slice (see Supplementary Figure S4). Evoked extracellular DA concentration ([DA]_o_) was measured using FSCV at CFMs as described previously (Threlfell et al., 2012). In brief, a triangular voltage waveform was scanned across the microelectrode (−700 to +1300 mV and back vs Ag/AgCl reference, scan rate 800 V/s) using a Millar Voltammeter (Julian Millar, Barts and the London School of Medicine and Dentistry), with a sweep frequency of 8 Hz. Electrical or light stimuli were delivered to the striatal slices at 2.5 min intervals, which allow stable release to be sustained at ∼90-95% (see Fig. 1B,D) over the time course of control experiments. Evoked currents were confirmed as DA by comparison of the voltammogram with that produced during calibration with applied DA in aCSF (oxidation peak +500-600 mV and reduction peak −200 mV). Currents at the oxidation peak potential were measured from the baseline of each voltammogram and plotted against time to provide profiles of [DA]_o_ versus time. CFMs were calibrated *post hoc* in 2 μM DA in each experimental solution. Calibration solutions were made immediately before use from stock solution of 2.5 mM DA in 0.1 M HClO_4_ stored at 4 °C. CFM sensitivity to DA was between 10 and 40 nA/μM. Unless noted otherwise, FSCV recordings were carried out in the presence of dihydro-β-erythroidine (DHβE, 1 μM), an antagonist at β2 subunit-containing nicotinic acetylcholine receptors (nAChRs), to eliminate cholinergic signalling effects on DA release (Exley and Cragg, 2008; Rice and Cragg, 2004; Threlfell et al., 2012). Release was tetrodotoxin-sensitive as shown previously (Threlfell et al., 2012).

In experiments where [DA]_o_ was evoked by electrical stimulation, a local bipolar concentric Pt/Ir electrode (25 μm diameter; FHC Inc.) was placed approximately 100 μm from the CFMs and stimulus pulses (200 μs duration) were given at 0.6 mA (perimaximal in drug-free control conditions). We applied either single pulses (1p) or 2-10 pulses (2p, 4p, 5p, 10p) at 10 - 100 Hz. A frequency of 100 Hz is useful as a tool for exposing changes in short-term plasticity in DA release that arise through changes in initial release probability (Jennings et al., 2015; Rice and Cragg, 2004). In experiments where [DA]_o_ was evoked by light stimulation in slices prepared from Slc6a3^*IRES-Cre*^ mice expressing ChR2, DA axons in striatum were activated by TTL-driven (Multi Channel Stimulus II, Multi Channel Systems) brief pulses (2 ms) of blue light (470 nm; 5 mWmm^-2^; OptoLED; Cairn Research), which illuminated the field of view (2.2 mm, x10 water-immersion objective). Epifluorescence used to visualize ChR2-eYFP expression was used sparingly to minimize ChR2 activation before recordings

### Electrophysiology

Individual slices were hemisected and transferred to a recording chamber and superfused at ∼3.0 mL/min with aCSF at 31-33 °C. Cells were visualized through a X40 water-immersion objective with differential interference contrast optics. All whole-cell experiments were recorded using borosilicate glass pipettes with resistances in the 3 – 5 MΩ range and were pulled on a Flaming-Brown micropipette puller (P-1000, Sutter Instruments). Whole-cell voltage-clamp electrophysiology recordings were made from spiny projection neurons (SPNs; identified by their membrane properties (Gertler et al., 2008; Planert et al., 2013)) in the DLS or NAcC. SPNs were voltage-clamped at −70 mV using a MultiClamp 700B amplifier (Molecular Devices) and with pipettes filled with a CsCl-based internal solution (in mM 120 CsCl, 15 CsMeSO_3_, 8 NaCl, 0.5 EGTA, 10 HEPES, 2 Mg-ATP, 0.3 Na-GTP, 5 QX-314; pH 7.3 adjusted with CsOH; osmolarity ranging from 305 - 310 mOsmkg^-1^). The recording perfusate always contained NBQX (5 μM) and APV (50 μM) to block AMPA and NMDA receptor-mediated inward currents. Errors due to the voltage drop across the series resistance (<20 MΩ) were left uncompensated and membrane potentials were corrected for a ∼5 mV liquid junction potential. Cells were discarded from analysis if if series resistance varied by more than 15% or increased over 25 MΩ.

To record tonic GABA_A_ currents, SPNs voltage-clamped at −70 mV were recorded in gap-free mode. Cells were allowed to stabilize for 5-10 min before drug manipulations: GAT inhibitors were bath applied for 20 −25 min; picrotoxin (100 μM) for an additional 3-5 min. Recordings of light-evoked GABA currents in SPNs from ChR2-expressing DA axons in slices from Slc6a3^*IRES-Cre*^ mice were taken 10 min after break-in, and at 30 s intervals for a duration of 10 min from SPNs voltage-clamped at −70 mV. Under these conditions, GABA_A_ receptor-mediated currents appear inward as reported previously (Tritsch et al., 2012). TTL-driven (Multi Channel Stimulus II, Multi Channel Systems) brief pulses (2 ms) of blue light (470 nm; 5 mWmm^-2^; OptoLED; Cairn Research) illuminated the full field of view (2.2 mm, X10 water-immersion objective).

### High-performance liquid chromatography

DA content in dorsal striatum was measured by HPLC with electrochemical detection as described previously (Janezic et al., 2013). Tissue punches (2 mm in diameter) were taken from dorsal striatum in two brain slices per animal, snap frozen and stored at −80 °C in 200 μL 0.1 M HClO_4_. On the day of analysis, samples were thawed on ice, homogenized, and centrifuged at 15,000 g for 15 min at 4 °C. The supernatant was analysed for DA content. Analytes were separated using a 4.6 × 250 mm Microsorb C18 reverse-phase column (Varian or Agilent) and detected using a Decade II SDS electrochemical detector with a Glassy carbon working electrode (Antec Leyden) set at + 0.7 V with respect to a Ag/AgCl reference electrode. The mobile phase consisted of 13% methanol (vol/vol), 0.12 M NaH_2_PO_4_, 0.5–4.0 mM octenyl succinic anhydride (OSA), and 0.8 mM EDTA (pH 4.4–4.6), and the flow rate was fixed at 1 mL/min.

### Western blot

Mouse brains were extracted and sliced using the procedures outlined above. One 1.2 mm thick coronal slice containing striatum was prepared from each brain and one tissue punch (2 mm in diameter) of dorsal striatum taken per hemisphere. Striatal tissue samples were snap-frozen and stored at at −80 °C. For analysis, striatal tissue was defrosted on ice, homogenized in RIPA Lysis and Extraction Buffer (Sigma) containing 150 mM NaCl, 1.0% IGEPAL, 0.5% sodium deoxycholate, 0.1% SDS, 50 mM Tris, pH 8.0, with Complete-Mini Protease Inhibitor and PhosStop (Roche), using a Tissue Tearor (Biospec Products, Inc), and soluble fraction isolated by microcentrifugation at 15,000 g for 15 min at 4°C. Total protein content was quantified using a BCA Protein Assay Kit (Thermo Scientific) and equal amounts of total protein were loaded onto 4 – 15% Tris-Glycine gels (BioRad). Following electrophoresis (200 V for ∼45 min), proteins were transferred onto polyvinylidene fluoride membranes (BioRad). Blots were probed overnight at 4°C with 1:1,000 rabbit anti-GABA transporter 1 (Synaptic Systems, 274102) or 1:1,000 rabbit anti-GABA transporter 3 (Abcam, AB181783). Blots were incubated with HRP-conjugated secondary anitbodies at 1:3,000 for 1 h at room temperature and bands developed using ECL Prime Western Blotting Detection Reagent (GE Healthcare). Blots were subsequently incubated with 1:20,000 HRP-conjugated β-actin (Abcam, AB49900) for 1 h at room temperature and bands developed as above. Visualization and imaging of blots was performed with a ChemiDoc Imaging System (BioRad) and bands quantified using Image Lab Software (BioRad). Protein concentration for GAT-1 and GAT-3 were normalized to β-actin.

### Indirect immunofluorescence

Adult mice were anaesthetized with an overdose of pentobarbital and transcardially perfused with 20-50 mL of phosphate-buffered saline (PBS), followed by 30-100 mL of 4% paraformaldehyde (PFA) in 0.1 M phosphate buffer, pH 7.4. Brains were removed and post-fixed overnight in 4% PFA. Brains were embedded in agar (3-4%) and coronal sections (50 µm) were cut on a vibrating microtome (Leica VT1000S) and collected in a 1 in 4 series. Sections were stored in PBS with 0.05% sodium azide. Upon processing, sections were washed in PBS and then blocked for 1h in a solution of PBS TritonX (0.3%) with sodium azide (0.02%; PBS-Tx) containing 10% normal donkey serum (NDS). Sections were then incubated in primary antibodies overnight in PBS-Tx with 2% NDS at 4°C. Primary antibodies: rabbit anti-TH (1:2,000, Sigma-Aldrich, ab112); rabbit anti-GAT1 (1:1,000, Synaptic Systems, 274102); rabbit anti-GAT3 (1:250, Millipore/Chemicon, AB1574); rabbi anti-NeuN (1:500, Biosensis, R-3770-100); guinea pig anti-S100β (1:2,000, Synaptic Systems, 287004); rat anti-GFP that also recognizes eYFP (1:1,000, Nacalai Tesque, 04404-84); and guinea-pig anti-parvalbumin (1:1,000, Synaptic Systems, 195004). Sections were then incubated in species-appropriate fluorescent secondary antibodies with minimal cross-reactivity overnight in PBS-Tx at room temperature (Donkey anti-Rabbit AlexaFluor 488, 1:1,000, Invitrogen, A21206; Donkey anti-Rabbit Cy3, 1:1,000, Jackson ImmunoResearch, 711-165-152; Donkey anti-Guinea Pig AlexaFluor 488, 1:1,000, Jackson ImmunoResearch, 706-545-148; Donkey anti-Rat AlexaFluor 488, 1:1,000, Jackson ImmunoResearch, 712-545-153). Sections were washed in PBS and then mounted on glass slides and cover-slipped using Vectashield (Vector Labs). Coverslips were sealed using nail varnish and stored at 4°C. To verify the specificity of ChR2-eYFP expression in TH-positive structures in Slc6a3^*IRES-Cre*^ mice, mounted sections were imaged with an Olympus BX41 microscope with Olympus UC30 camera and filters for appropriate excitation and emission wave lengths (Olympus Medical).

### Confocal imaging and image analysis

Confocal images were acquired with an LSM880/Axio.Imager Z2 (Zeiss) and Image J was used for image analysis. For whole striatum analysis of GAT1 or GAT3, the X10 (NA = 0.45) objective was used and all imaging settings (laser %, pinhole/optical section, pixel size, gain, and scanning speed) were kept constant between animals. For the quantification of fluorescence (mean grey values), 4 sections in the rostro-caudal plane were imaged at approximately the following distances rostral of Bregma; +1.3mm, +1.0mm, +0.6mm, and +0.25mm (see Supplementary Figure S4). A region of interest (ROI) of 300 µm x 300 µm was overlaid over the DLS and the ventral CPu (vCPu); and an ROI of 200 µm X 200 µm was overlaid on NAcC and the NAcS, for both hemispheres. Values for NAcC and NAcS were taken from the 2 most rostral sections (see Supplementary Figure 4). Mean grey values from the areas of interest were normalized to the median grey value for each hemisphere (n = 12 hemispheres from 6 animals). For examination of co-localization a X63 objective was used (NA = 1.46); Z-stacks were taken with the pinhole set to 1 Airy Unit (optical section = 0.7µm) with a z-stack interval of 0.30 µm or 0.35 µm. In order to assess co-localization ZEN (blue edition v.2.3; Zeiss) software was used. For S100β, PV+ axons (eYFP in PV-Cre mice) or DA axons (eYFP in *Slc6a3*^*IRES-Cre*^ mice) and GAT1/3 co-localization, stacks from a minimum of 2 striatal regions and 2 NAcC regions in at least one section were examined per animal (n = 3 per marker).

### Drugs

(S)-SNAP5114 (SNAP, 50 µM), (±)-nipecotic acid (NPA, 1.5 mM), γ-aminobutyric acid (GABA, 2 mM) and picrotoxin (100 μM) were obtained from Sigma Aldrich. Dihydro-β-erythroidine hydrobromide (DHβE, 1μM), (+)-bicuculline (10 μM) and tetrodotoxin (1 μM) were obtained from Tocris Bioscience. DL-2-Amino-5-phosphonovaleric acid (AP5, 50 μM), disulfiram (10 μM) and SKF-89976A hydrochloride (SKF, 20 μM) were obtained from Santa Cruz Biotechnology. NBQX disodium salt (NBQX, 5 μM) and CGP 55845 hydrochloride (CGP, 4 μM) were obtained from Abcam. Fluorocitrate was prepared as previously described (Paulsen et al., 1987). In brief, D,L-fluorocitric acid Ba_3_ salt (Sigma Aldrich) was dissolved in 0.1 M HCl, the Ba^2+^ precipitated with 0.1 M Na_2_S0_4_ and then centrifuged at 1,000 g for 5 min. Supernatant containing fluorocitrate was used at a final concentration of 200 µM for experimentation. All drugs were dissolved in distilled water or dimethyl sulfoxide (DMSO) to make stock aliquots at 1,000–10,000 × final concentrations and stored at −20 °C. Stock aliquots were diluted with aCSF to final concentration immediately before use.

### Data acquisition and analysis

FSCV data were digitized at 50 kHz using a Digidata 1550A digitizer (Molecular Devices). Data were acquired and analyzed using Axoscope 11.0 (Molecular Devices) and locally written VBA scripts. For drug effects, peak [DA]_o_ was averaged over 4 stimulations once peak [DA]_o_ had re-stabilized post-drug application and compared to time-matched data from drug-free controls, unless otherwise stated. Electrically evoked [DA]_o_ in slices pre-incubated in fluorocitrate (200 µM) exhibited modest run-down across repeated 1p stimulations (n = 5 experiments/3 mice; data not shown) and therefore we used an alternative stimulation paradigm to compare a large number of dorsal striatal recording sites in slices pre-treated with fluorocitrate versus control conditions to minimize run-down. There are previous reports of a fluorocitrate-dependent slow rundown of excitatory postsynaptic currents in the hippocampus (Boddum et al., 2016), possibly reflecting astroglia’s role in stable synaptic neurotransmission.

FSCV data are normalized to pre-drug conditions for clarity and for comparisons between regions. For experiments involving multiple pulse protocols, each stimulation type was repeated in triplicate, interspersed with 1p stimulations, and then averaged and normalized to 1p stimulations at each recording site, as previously (Lopes et al., 2019; Threlfell et al., 2012).

Membrane currents from voltage-clamp electrophysiology experiments were amplified and low-pass filtered at 5 kHz using a MultiClamp 700B amplifier (Molecular Devices), digitized at 10 kHz and acquired using a Digidata 1550A digitizer (Molecular Devices). Peak amplitude, onset latency, peak latency, 10-90% rise time and decay time were measured from an average of 3 replicate traces recorded before and after drug wash on conditions using Clampfit 10.4.1.4 software (Molecular Devices).

For all experiments, data were collected from a minimum of 3 animals. Data were compared for statistical significance using Prism 7 (Graph Pad) with the following statistical tests (as indicated in the text, and two-tailed): un-paired t-tests, paired t-tests, two-way repeated-measures ANOVA followed by Sidak’s multiple comparison tests, and where the data were not normally distributed, Mann-Whitney U tests, Kruskal-Wallis ANOVA followed by Dunn’s Multiple Comparisons, Friedman’s ANOVA on Ranks and Student-Newman-Keuls multiple comparisons and for comparing cumulative distributions, Komogorov-Smirnov tests. p values smaller than 0.05 were considered statistically significant, adjusted for multiple comparisons.

## ACKNOWLEDGEMENTS

This work is supported by grants from Parkinson’s UK (G-1504, and Monument Trust Discovery Award J-1403), a Clarendon Fund Studentship awarded to B.M.R. and a Fundação para a Ciência e a Tecnologia studentship awarded to E.F.L. N.M.D. and P.J.M. were supported by the UK Medical Research Council (Award MC_UU_12024/2 to P.J.M.) and the Wellcome Trust (Investigator Award 101821 to P.J.M.). We thank Ben Micklem and Lisa Conyers for their assistance with anatomical work, Milena Cioroch for technical expertise in maintaining transgenic mouse colonies, and members of the Cragg laboratory for discussions throughout the course of this study. We thank Drs Yoland Smith and Jean-Francois Pare for GAT antibody recommendations.

## AUTHOR CONTRIBUTIONS

B.M.R. and S.J.C. conceived and designed the research, and wrote the manuscript with input from all other authors. B.M.R. performed the majority of experiments and analysis of data. N.M.D. and P.J.M. collected, analysed and interpreted anatomical confocal imaging data. K.R.B, E.F.L., S.T. and R.E.S. assisted with the collection, analysis and interpretation of voltammetry data. N.C.-R. and N.B.-V. provided expertise and assistance with western blot experiments. R.W.-M. conceived of and provided *SNCA*-OVX mice and assisted in interpretation of data.

## REFERENCES

Ade, K.K., Janssen, M.J., Ortinski, P.I., and Vicini, S. (2008). Differential Tonic GABA Conductances in Striatal Medium Spiny Neurons. J. Neurosci. 28, 1185–1197.

Augood, S.J., Herbison, A.E., and Emson, P.C. (1995). Localization of GAT-1 GABA transporter mRNA in rat striatum: cellular coexpression with GAD67 mRNA, GAD67 immunoreactivity, and parvalbumin mRNA. J. Neurosci. 15, 865–874.

Boddum, K., Jensen, T.P., Magloire, V., Kristiansen, U., Rusakov, D.A., Pavlov, I., and Walker, M.C. (2016). Astrocytic GABA transporter activity modulates excitatory neurotransmission. Nat. Commun. 7, 13572.

Bonansco, C., Couve, A., Perea, G., Ferradas, C.A., Roncagliolo, M., and Fuenzalida, M. (2011). Glutamate released spontaneously from astrocytes sets the threshold for synaptic plasticity. Eur. J. Neurosci. 33, 1483–1492.

Borden, L.A. (1996). GABA transporter heterogeneity: Pharmacology and cellular localization. Neurochem. Int. 29, 335–356.

Brickley, S.G., and Mody, I. (2012). Extrasynaptic GABAA Receptors: Their Function in the CNS and Implications for Disease. Neuron 73, 23–34.

Brimblecombe, K.R., Gracie, C.J., Platt, N.J., and Cragg, S.J. (2015). Gating of dopamine transmission by calcium and axonal N-, Q-, T- and L-type voltage-gated calcium channels differs between striatal domains. J. Physiol. 593, 929–946.

Britt, J.P., and McGehee, D.S. (2008). Presynaptic Opioid and Nicotinic Receptor Modulation of Dopamine Overflow in the Nucleus Accumbens. J. Neurosci. 28, 1672–1681.

Cepeda, C., Galvan, L., Holley, S.M., Rao, S.P., Andre, V.M., Botelho, E.P., Chen, J.Y., Watson, J.B., Deisseroth, K., and Levine, M.S. (2013). Multiple Sources of Striatal Inhibition Are Differentially Affected in Huntington’s Disease Mouse Models. J. Neurosci. 33, 7393–7406.

Chai, H., Diaz-Castro, B., Shigetomi, E., Monte, E., Octeau, J.C., Yu, X., Cohn, W., Rajendran, P.S., Vondriska, T.M., Whitelegge, J.P., et al. (2017). Neural Circuit-Specialized Astrocytes: Transcriptomic, Proteomic, Morphological, and Functional Evidence. Neuron 95, 531-549.e9.

Charara, A., Heilman, C., Levey, A.., and Smith, Y. (1999). Pre- and postsynaptic localization of GABAB receptors in the basal ganglia in monkeys. Neuroscience 95, 127–140.

Chazalon, M., Paredes-Rodriguez, E., Morin, S., Martinez, A., Cristóvão-Ferreira, S., Vaz, S., Sebastiao, A., Panatier, A., Boué-Grabot, E., Miguelez, C., et al. (2018). GAT-3 Dysfunction Generates Tonic Inhibition in External Globus Pallidus Neurons in Parkinsonian Rodents. Cell Rep. 23, 1678–1690.

Dodson, P.D., Dreyer, J.K., Jennings, K.A., Syed, E.C.J., Wade-Martins, R., Cragg, S.J., Bolam, J.P., and Magill, P.J. (2016). Representation of spontaneous movement by dopaminergic neurons is cell-type selective and disrupted in parkinsonism. Proc. Natl. Acad. Sci. U. S. A. 113, E2180–8.

Durkin, M.M., Smith, K.E., Borden, L.A., Weinshank, R.L., Branchek, T.A., and Gustafson, E.L. (1995). Localization of messenger RNAs encoding three GABA transporters in rat brain: an in situ hybridization study. Mol. Brain Res. 33, 7–21.

Exley, R., and Cragg, S.J. (2008). Presynaptic nicotinic receptors: A dynamic and diverse cholinergic filter of striatal dopamine neurotransmission. In British Journal of Pharmacology, (John Wiley & Sons, Ltd (10.1111)), pp. S283–S297.

Farrant, M., and Nusser, Z. (2005). Variations on an inhibitory theme: phasic and tonic activation of GABAA receptors. Nat. Rev. Neurosci. 6, 215–229.

Ficková, M., Dahmen, N., Fehr, C., and Hiemke, C. (1999). Quantitation of GABA transporter 3 (GAT3) mRNA in rat brain by competitive RT-PCR. Brain Res. Protoc. 4, 341–350.

Gertler, T.S., Chan, C.S., and Surmeier, D.J. (2008). Dichotomous Anatomical Properties of Adult Striatal Medium Spiny Neurons. J. Neurosci. 28, 10814–10824.

Gokce, O., Neff, N.F., Fuccillo, M. V., Südhof, T.C., Treutlein, B., Quake, S.R., Malenka, R.C., Stanley, G.M., Rothwell, P.E., and Camp, J.G. (2016). Cellular Taxonomy of the Mouse Striatum as Revealed by Single-Cell RNA-Seq. Cell Rep. 16, 1126–1137.

Goubard, V., Fino, E., and Venance, L. (2011). Contribution of astrocytic glutamate and GABA uptake to corticostriatal information processing. J. Physiol. 589, 2301–2319.

Henneberger, C., Papouin, T., Oliet, S.H.R., and Rusakov, D.A. (2010). Long-term potentiation depends on release of d-serine from astrocytes. Nature 463, 232–236.

Janezic, S., Threlfell, S., Dodson, P.D., Dowie, M.J., Taylor, T.N., Potgieter, D., Parkkinen, L., Senior, S.L., Anwar, S., Ryan, B., et al. (2013). Deficits in dopaminergic transmission precede neuron loss and dysfunction in a new Parkinson model. Proc. Natl. Acad. Sci. 110, E4016–E4025.

Jennings, K.A., Platt, N.J., and Cragg, S.J. (2015). The impact of a parkinsonian lesion on dynamic striatal dopamine transmission depends on nicotinic receptor activation. Neurobiol. Dis. 82, 262–268.

Jin, X.-T., Galvan, A., Wichmann, T., and Smith, Y. (2011). Localization and Function of GABA Transporters GAT-1 and GAT-3 in the Basal Ganglia. Front. Syst. Neurosci. 5, 1–10.

Kaeser, P.S., and Regehr, W.G. (2013). Molecular Mechanisms for Synchronous, Asynchronous, and Spontaneous Neurotransmitter Release. Annu. Rev. Physiol. 76, 333–363.

Kim, J.I., Ganesan, S., Luo, S.X., Wu, Y.W., Park, E., Huang, E.J., Chen, L., and Ding, J.B. (2015). Aldehyde dehydrogenase 1a1 mediates a GABA synthesis pathway in midbrain dopaminergic neurons. Science (80-.). 350, 102–106.

Kirmse, K., Dvorzhak, A., Kirischuk, S., and Grantyn, R. (2008). GABA transporter 1 tunes GABAergic synaptic transmission at output neurons of the mouse neostriatum. J. Physiol. 586, 5665–5678.

Kirmse, K., Kirischuk, S., and Grantyn, R. (2009). Role of GABA transporter 3 in GABAergic synaptic transmission at striatal output neurons. Synapse 63, 921–929.

Li, G., Shao, C., Chen, Q., Wang, Q., and Yang, K. (2017). Accumulated GABA activates presynaptic GABAB receptors and inhibits both excitatory and inhibitory synaptic transmission in rat midbrain periaqueductal gray. Neuroreport 28, 313–318.

Liu, J., Zhou, Z., Hodgkinson, C.A., Yuan, Q., Shen, P.-H., Mulligan, C.J., Wang, A., Gray, R.R., Roy, A., Virkkunen, M., et al. (2011). Haplotype-Based Study of the Association of Alcohol-Metabolizing Genes With Alcohol Dependence in Four Independent Populations. Alcohol. Clin. Exp. Res. 35, 304–316.

Lopes, E.F., Roberts, B.M., Siddorn, R.E., Clements, M.A., and Cragg, S.J. (2019). Inhibition of nigrostriatal dopamine release by striatal GABAA and GABAB receptors. J. Neurosci. 2028–18.

Lozovaya, N., Eftekhari, S., Cloarec, R., Gouty-Colomer, L.A., Dufour, A., Riffault, B., Billon-Grand, M., Pons-Bennaceur, A., Oumar, N., Burnashev, N., et al. (2018). GABAergic inhibition in dual-transmission cholinergic and GABAergic striatal interneurons is abolished in Parkinson disease. Nat. Commun. 9, 1422.

Martín, E.D., Fernández, M., Perea, G., Pascual, O., Haydon, P.G., Araque, A., and Ceña, V. (2007). Adenosine released by astrocytes contributes to hypoxia-induced modulation of synaptic transmission. Glia 55, 36–45.

Matsuda, W., Furuta, T., Nakamura, K.C., Hioki, H., Fujiyama, F., Arai, R., and Kaneko, T. (2009). Single Nigrostriatal Dopaminergic Neurons Form Widely Spread and Highly Dense Axonal Arborizations in the Neostriatum. J. Neurosci. 29, 444–453.

Ng, C.H., Wang, X.S., and Ong, W.Y. (2000). A light and electron microscopic study of the GABA transporter GAT-3 in the monkey basal ganglia and brainstem. J. Neurocytol. 29, 595–603.

Oorschot, D.E. (1996). Total number of neurons in the neostriatal, pallidal, subthalamic, and substantia nigral nuclei of the rat basal ganglia: A stereological study using the cavalieri and optical disector methods. J. Comp. Neurol. 366, 580–599.

Paulsen, R.E., Contestabile, A., Villani, L., and Fonnum, F. (1987). An In Vivo Model for Studying Function of Brain Tissue Temporarily Devoid of Glial Cell Metabolism: The Use of Fluorocitrate. J. Neurochem. 48, 1377–1385.

Pitman, K.A., Puil, E., and Borgland, S.L. (2014). GABAB modulation of dopamine release in the nucleus accumbens core. Eur. J. Neurosci. 40, 3472–3480.

Planert, H., Berger, T.K., and Silberberg, G. (2013). Membrane Properties of Striatal Direct and Indirect Pathway Neurons in Mouse and Rat Slices and Their Modulation by Dopamine. PLoS One 8, e57054.

Rice, M.E., and Cragg, S.J. (2004). Nicotine amplifies reward-related dopamine signals in striatum. Nat. Neurosci. 7, 583–584.

Santhakumar, V., Jones, R.T., and Mody, I. (2010). Developmental regulation and neuroprotective effects of striatal tonic GABAA currents. Neuroscience 167, 644–655.

Schmitz, Y., Schmauss, C., and Sulzer, D. (2002). Altered Dopamine Release and Uptake Kinetics in Mice Lacking D 2 Receptors. J. Neurosci. 22, 8002–8009.

Schmitz, Y., Benoit-Marand, M., Gonon, F., and Sulzer, D. (2003). Presynaptic regulation of dopaminergic neurotransmission. J. Neurochem. 87, 273–289.

Sherva, R., Rice, J.P., Neuman, R.J., Rochberg, N., Saccone, N.L., and Bierut, L.J. (2009). Associations and Interactions Between SNPs in the Alcohol Metabolizing Genes and Alcoholism Phenotypes in European Americans. Alcohol. Clin. Exp. Res. 33, 848–857.

Shin, J.H., Adrover, M.F., and Alvarez, V.A. (2017). Distinctive Modulation of Dopamine Release in the Nucleus Accumbens Shell Mediated by Dopamine and Acetylcholine Receptors. J. Neurosci. 37, 11166–11180.

Sloan, M., Alegre-Abarrategui, J., Potgieter, D., Kaufmann, A.-K., Exley, R., Deltheil, T., Threlfell, S., Connor-Robson, N., Brimblecombe, K., Wallings, R., et al. (2016). *LRRK2* BAC transgenic rats develop progressive, L-DOPA-responsive motor impairment, and deficits in dopamine circuit function. Hum. Mol. Genet. 25, 951–963.

Sulzer, D., Cragg, S.J., and Rice, M.E. (2016). Striatal dopamine neurotransmission: Regulation of release and uptake. Basal Ganglia.

Taylor, T.N., Potgieter, D., Anwar, S., Senior, S.L., Janezic, S., Threlfell, S., Ryan, B., Parkkinen, L., Deltheil, T., Cioroch, M., et al. (2014). Region-specific deficits in dopamine, but not norepinephrine, signaling in a novel A30P α-synuclein BAC transgenic mouse. Neurobiol. Dis. 62, 193–207.

Threlfell, S., and Cragg, S.J. (2011). Dopamine Signaling in Dorsal Versus Ventral Striatum: The Dynamic Role of Cholinergic Interneurons. Front. Syst. Neurosci. 5, 11.

Threlfell, S., Clements, M.A., Khodai, T., Pienaar, I.S., Exley, R., Wess, J., and Cragg, S.J. (2010). Striatal Muscarinic Receptors Promote Activity Dependence of Dopamine Transmission via Distinct Receptor Subtypes on Cholinergic Interneurons in Ventral versus Dorsal Striatum.

Threlfell, S., Lalic, T., Platt, N.J., Jennings, K.A., Deisseroth, K., and Cragg, S.J. (2012). Striatal dopamine release is triggered by synchronized activity in cholinergic interneurons. Neuron 75, 58–64.

Tritsch, N.X., Ding, J.B., and Sabatini, B.L. (2012). Dopaminergic neurons inhibit striatal output through non-canonical release of GABA. Nature 490, 262–266.

Tritsch, N.X., Oh, W.J., Gu, C., and Sabatini, B.L. (2014). Midbrain dopamine neurons sustain inhibitory transmission using plasma membrane uptake of GABA, not synthesis. Elife 3, e01936.

Wójtowicz, A.M., Dvorzhak, A., Semtner, M., and Grantyn, R. (2013). Reduced tonic inhibition in striatal output neurons from Huntington mice due to loss of astrocytic GABA release through GAT-3. Front. Neural Circuits 7, 188.

Yasumi, M., Sato, K., Shimada, S., Nishimura, M., and Tohyama, M. (1997). Regional distribution of GABA transporter 1 (GAT1) mRNA in the rat brain: comparison with glutamic acid decarboxylase67 (GAD67) mRNA localization. Brain Res. Mol. Brain Res. 44, 205–218.

Yu, X., Taylor, A.M.W., Nagai, J., Golshani, P., Evans, C.J., Coppola, G., and Khakh, B.S. (2018). Reducing Astrocyte Calcium Signaling In Vivo Alters Striatal Microcircuits and Causes Repetitive Behavior. Neuron 99, 1170-1187.e9.

Zaccara, G., Cincotta, M., Borgheresi, A., and Balestrieri, F. (2004). Adverse motor effects induced by antiepileptic drugs.

Zhang, Y., Chen, K., Sloan, S.A., Bennett, M.L., Scholze, A.R., O’Keeffe, S., Phatnani, H.P., Guarnieri, P., Caneda, C., Ruderisch, N., et al. (2014). An RNA-Sequencing Transcriptome and Splicing Database of Glia, Neurons, and Vascular Cells of the Cerebral Cortex. J. Neurosci. 34, 11929–11947.

Zott, B., Simon, M.M., Hong, W., Unger, F., Chen-Engerer, H.J., Frosch, M.P., Sakmann, B., Walsh, D.M., and Konnerth, A. (2019). A vicious cycle of β amyloid–dependent neuronal hyperactivation. Science (80-.). 365, 559–565.

