## Supplementary Data for "Astrocytic striatal GABA transporter activity governs dopamine release and shows maladaptive downregulation in early parkinsonism"

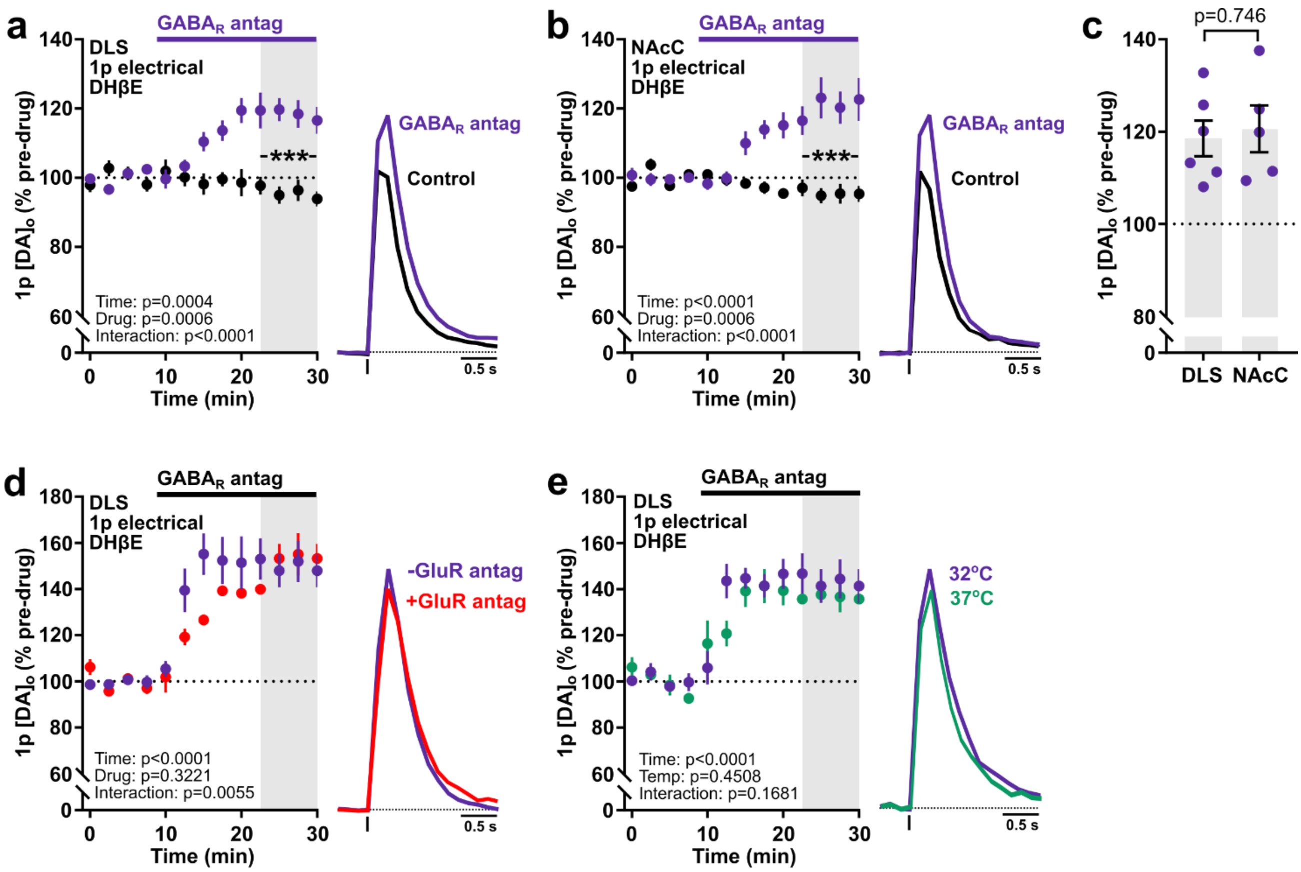


**Supplementary Fig 1. Tonic inhibition of striatal DA release is independent of nAChRs, glutamate input and temperature.** (**a**-**b**) *Left:* Mean peak [DA]$_o$ (± SEM) during consecutive recordings evoked by a single electrical pulse (*1p*) in control conditions (*black*, n = 9 experiments/7 mice for DLS, n = 6 experiments/5 mice for NAcC) and with GABA$_A$ and GABA$_B$ receptor antagonists (*solid bar*) (*purple*, *GABA$_R$ antag*), (+)-bicuculline (10 µM) and CGP 55845 (4 µM), respectively, recorded in the DLS (a, n = 6 experiments/3 mice) or NAcC (b, n = 5 experiments/3 mice), in the presence of nAChR antagonist DHβE (1 µM). *Right:* mean transients of [DA]$_o$ (normalized to pre-drug baselines) from last 4 time points (gray shaded region). (**c**) Mean peak [DA]$_o$ (± SEM) evoked by 1p following GABA$_R$ antagonism in DLS and NAcC (as % of pre-drug baseline). (**d**-**e**) Mean peak [DA]$_o$ (± SEM) during consecutive recordings evoked by a single electrical pulse (1p) in DLS before and during bath application of GABA$_R$ antagonists in either the absence (*purple*, n = 5 experiments/3 mice) or presence (*red*, n = 5 experiments/3 mice, d) of antagonists of ionotropic and metabotropic glutamate receptors (GluRs) , D-APV (50 µM), NBQX (5 µM), MCPG (200 µM), or recorded at 32 °C (*purple*, n = 5 experiments/3 mice) or 37 °C (*green*, n = 4 experiments/3 mice). Data are normalized to first 4 time points (*dotted line*). Mean transients of [DA]$_o$ (normalized to pre-drug baselines) from last 4 time points (*gray shaded region*). Statistical significance was assessed with two-way repeated measures ANOVA with Sidak's multiple comparison tests (a-b, c-e) and unpaired Student's t-test (c). DHβE (1 µM) present throughout. ****p* < 0.001.

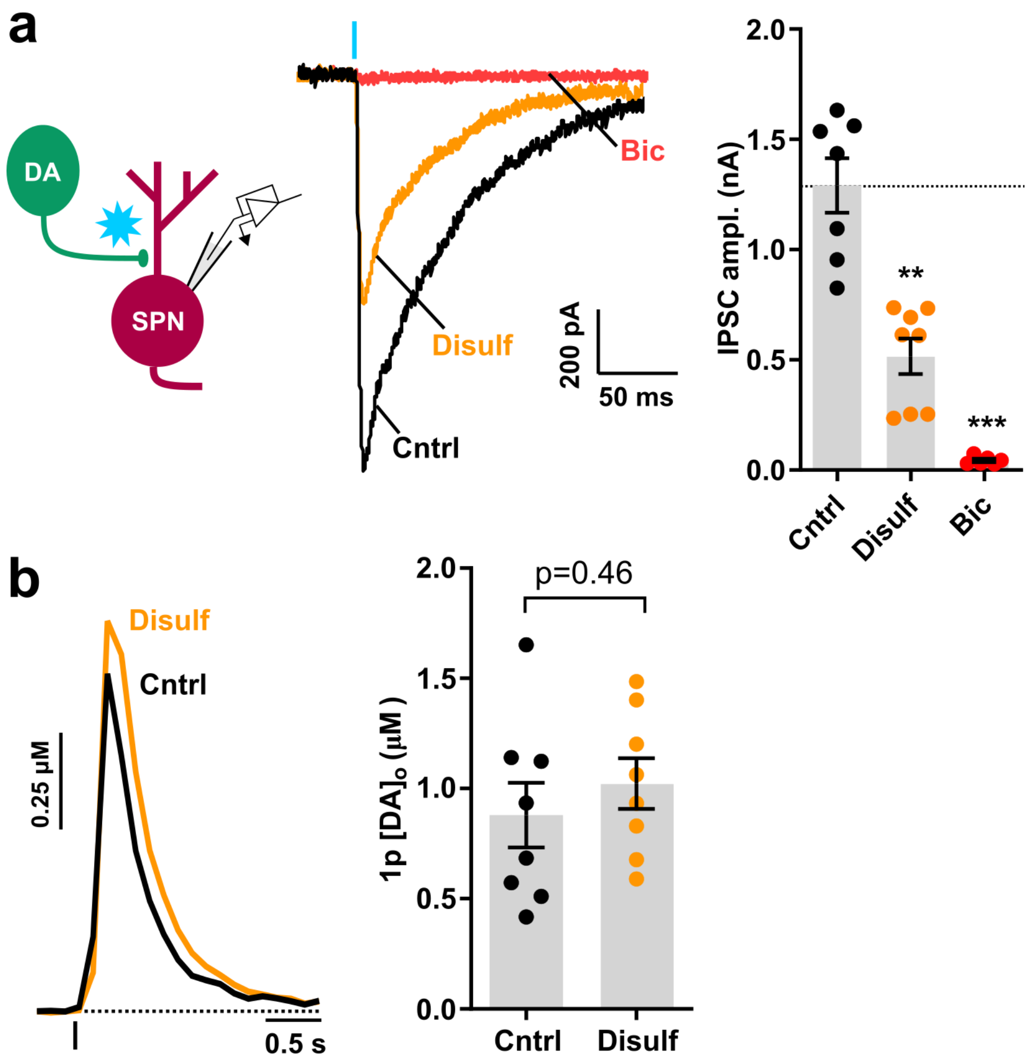


**Supplementary Fig 2. Inhibitory postsynaptic currents in SPNs resulting from GABA co-release from DA axons depend on ALDH.** (**a**) Representative and mean (± SEM) peak light-evoked inhibitory postsynaptic currents recorded from spiny projection neurons (SPNs) in the DLS of Slc6a3$^{IRES\text{-}Cre}$ mice expressing ChR2-eYFP in DA axons, voltage clamped at -70 mV and in the presence of ionotropic glutamate receptor antagonists (5 µM NBQX, 50 µM D-APV) in the perfusate. Light-evoked IPSCs were recorded in control conditions (*black*, n = 8 cells/5 mice) and in the presence of the $GABA_A$ receptor antagonist bicuculline (BIC, 10 µM) (*red*, n = 6 cells/4 mice), or the ALDH inhibitor disulfiram (Disulf, 10 µM) (*orange*, n = 8 cells/5 mice). Blue stimulus line indicates a 2 ms flash of blue light (470 nm LED, 5 mW×mm$^{-2}$ ). Control IPSCs had onset latencies of 2.27 ± 0.09 ms, thought to be consistent with monosynaptic transmission, and kinetics were slow (10 – 90% rise time: 4.39 ± 0.9 ms; 90 – 10 % decay time: 149 ± 13.8 ms), as previously described (Tritsch et al., 2012).
(**b**) ALDH inhibition with disulfiram (10 µM) did not alter $[DA]_o$ evoked by 1p electrical stimulation. Statistical significance was assessed by Mann-Whitney U tests in comparison to control. ***p* < 0.01, ****p* < 0.001.

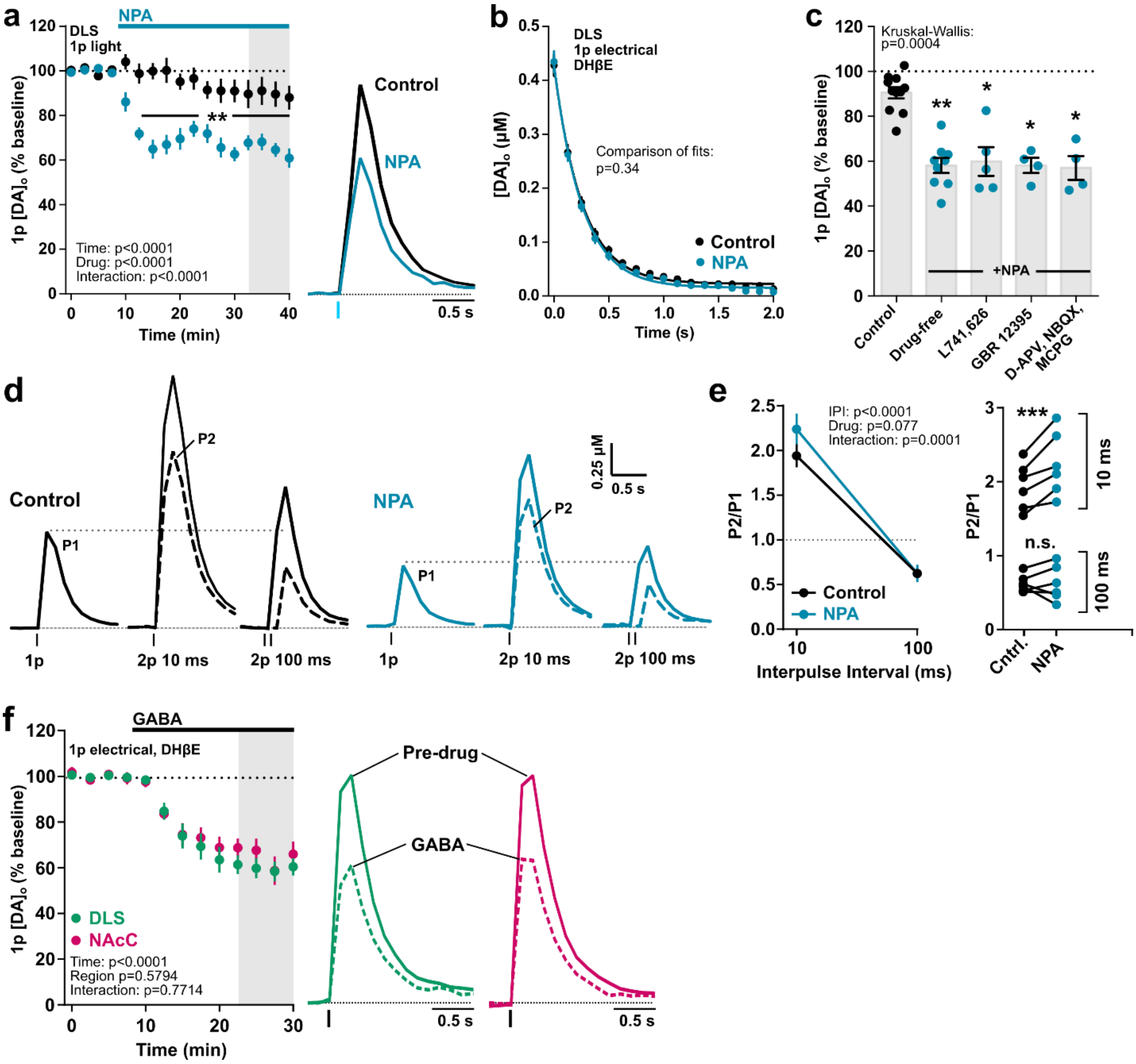


**Supplementary Fig 3. GAT inhibition attenuates DA release in DLS independently from mechanisms governing DA uptake, striatal $D_2$- dopamine receptors and glutamate receptors.**

(**a**) *Left*: Mean peak $[DA]_o$ (± SEM) during consecutive recordings evoked by a single light pulse in DLS in the absence (control, *black,* n = 8 experiments/6 mice) or presence of the non-specific GAT inhibitor (±)-nipecotic acid (NPA, 1.5 mM) (*blue*, n = 9 experiments/5 mice). Data are normalized to first 4 time points (*dotted line*). *Right*: mean transients of $[DA]_o$ (normalized to pre-drug baselines) from last 4 time points (*gray shaded region*). (**b**) Concentration-matched $[DA]_o$ (± SEM) decay over time in control conditions (*black*; $R^2$ = 0.96) or with the GAT inhibitor NPA (*blue*, $R^2$ = 0.96) from (Fig. 2a). (**c**) Mean peak $[DA]_o$ (± SEM) evoked by 1p following GAT inhibition (expressed as a % of pre-drug baseline) in control conditions (*black*) or following GAT inhibition with NPA (*blue*) in the absence (drug-free) or presence of $D_2$ antagonist L741,626 (2 µM) or DA transporter inhibitor GBR 12395 (1 µM) or ionotropic and metabotropic glutamate receptor inhibitors D-APV (50 µM), NBQX (5 µM) and MCPG (200 µM). Note the addition of these extra drugs did not alter NPA effect sizes. (**d**) Mean profiles of $[DA]_o$ transients elicited by single-pulse (1p) or paired-pulse (2p) electrical stimulation at inter-pulse intervals of 10 or 100 ms recorded in DLS in the absence (control, *black*, n = 6 experiments/4 mice) or presence of NPA (1.5 mM, *blue*, n = 6 experiments/ 4 mice). Solid traces exhibit $[DA]_o$ detected after stimulation (P1, P1 + P2), dotted traces show DA release attributed to the second pulse (P2) following subtraction of P1. DHβE (1 µM) was present throughout. (**e**) *Left*: Mean P2/P1 (± SEM) quantified from peak $[DA]_o$ shown in (a). *Right*: individual P2/P1 paired values. (**f**) *Left*: Mean peak $[DA]_o$ (± SEM) during consecutive recordings evoked by a single electrical pulse with bath application of GABA (2 mM) in DLS (*green*, n = 6 experiments/3 mice) or NAcC (*pink*, n = 5 experiments/3 mice). Data are normalized to first 4 time points (*dotted line*). *Right*: mean transients of $[DA]_o$ (normalized to pre-drug baselines) from last 4 time points (*gray shaded region*). DHβE (1 µM) present throughout. Statistical significance was assessed with two-way repeated measures ANOVA with Sidak's multiple comparison tests (a, e, f), extra sum-of-squares comparisons of fits (b), and Kruskal-Wallis test and Dunn's multiple comparisons to control conditions (c). **p* < 0.05, ***p* < 0.01, ****p* < 0.001.

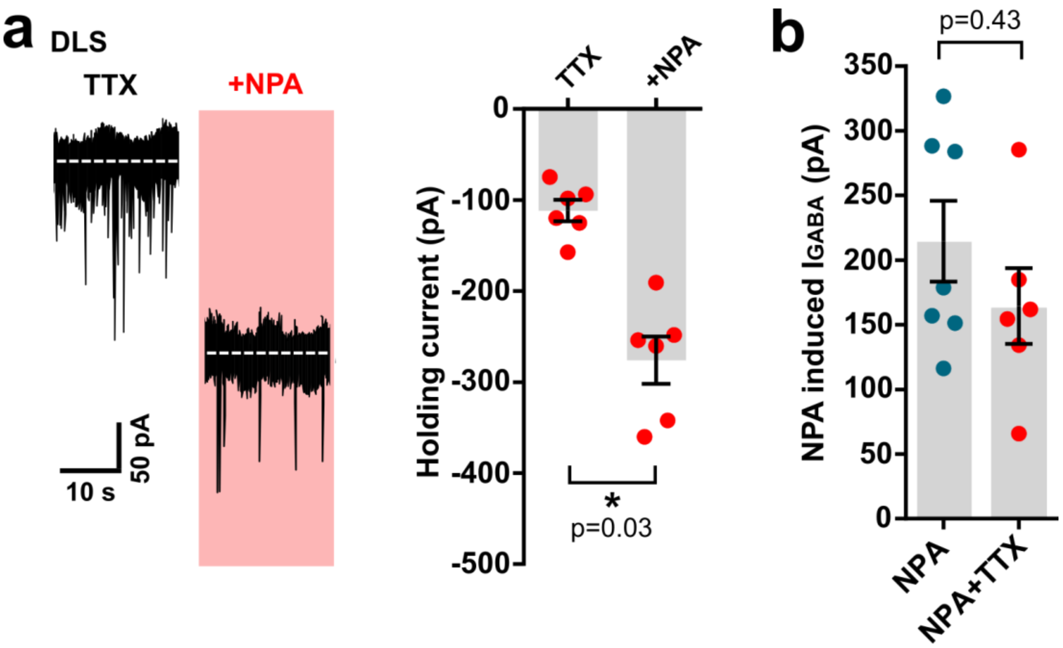


**Supplementary Fig 4. Tonic GABA currents in striatal projection neurons (SPNs) are augmented by GAT inhibition in an action potential-independent manner.** (**a**) *Left*, representative continuous whole-cell recordings from SPNs in DLS voltage-clamped at -70 mV in the presence of ionotropic glutamate receptor antagonists NBQX (5 μM) and D-AP5 (50 μM) and $Na_v$ blocker TTX (1 μM), before and during bath application of GAT inhibitor NPA (*red*, n = 6 cell/3 mice). NPA increase the extracellular $GABA_A$-mediated inward current, revealed by a shift in the holding current. *Right*, mean (± SEM) holding current in pA recorded in SPNs in control conditions and upon addition of NPA. (**b**) Mean (± SEM) tonic $GABA_A$-receptor-mediated currents induced by NPA recorded from SPNs in the absence (*blue*, n = 7 cells/5 mice, from main Fig. 4) or presence of TTX (*red*, n = 6 cells/3 mice) calculated by subtracting pre-drug holding current from GAT block-induced holding current. Statistical significance was assessed with Mann-Whitney U tests. *$p < 0.05$.

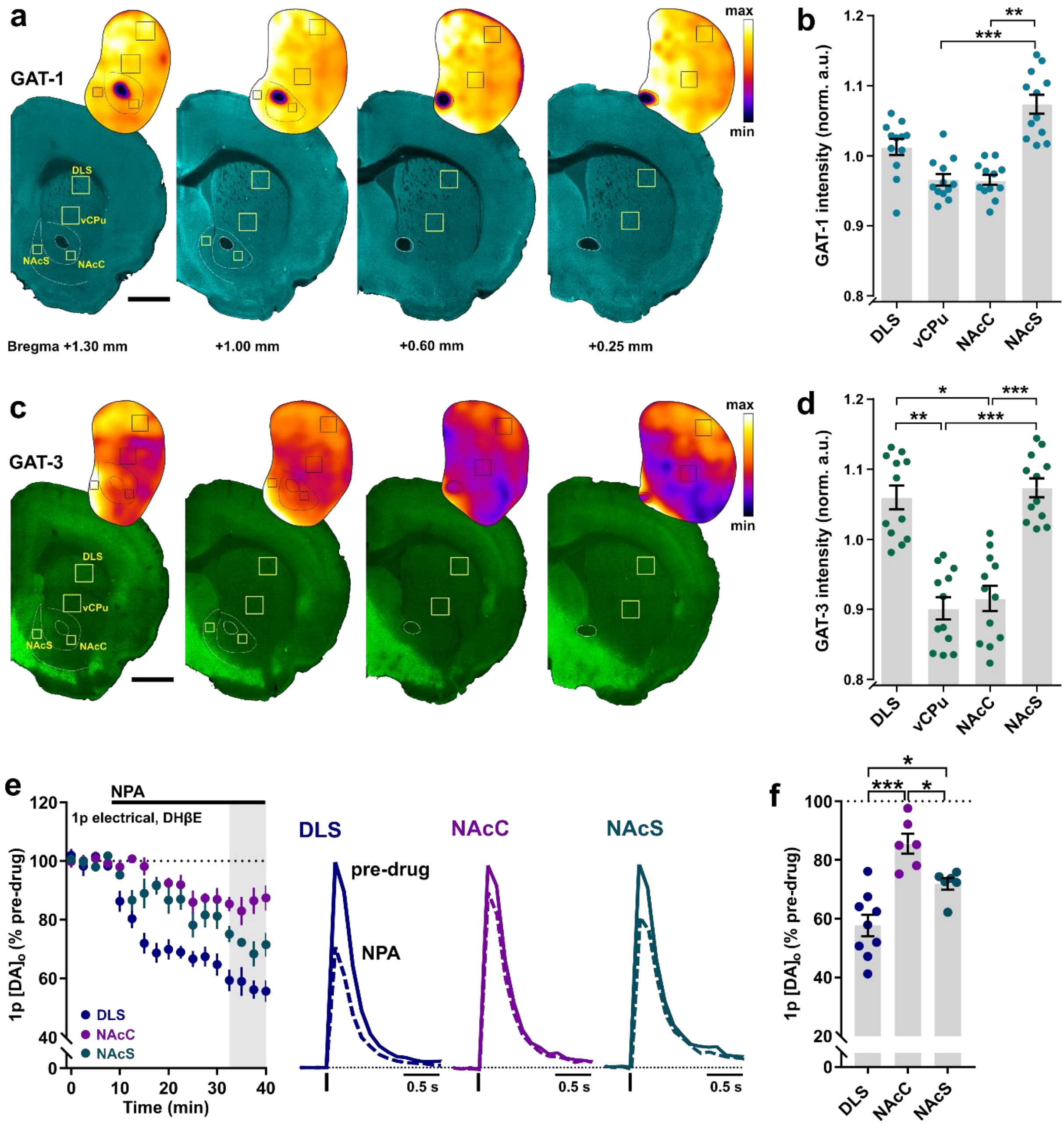


**Supplementary Fig 5. Heterogenous GAT-1 and GAT-3 expression and regulation of DA release across striatal territories.** (**a**, **c**) Representative immunofluorescence signals for GAT-1 (*cyan*, a) and GAT-3 (*green*, c) using confocal microscopy in coronal sections across the rostral-caudal limits containing striatum prepared from an individual C57BL/6J mouse with heat maps for striatal GAT intensity. Representative locations for GAT intensity measurements in the dorsolateral striatum (DLS), ventral CPu (vCPu), nucleus accumbens core (NAcC) and shell (NAcS) are indicated by boxes. Scale bars: 1 mm. (**b**, **d**) Mean (± SEM) GAT-1 (b) and GAT-3 (d) fluorescence intensities across striatal territories. Intensity (grey) values were normalized to the median of all striatal measurements within each hemisphere and then averaged across rostral-caudal sites (n = 12 hemispheres/6 mice for each GAT-1 and GAT-3). (**e**) Plot of consecutive recordings of mean peak $[DA]_o$ (± SEM) evoked by a single electrical pulse (1p) before and during the bath application of the nonspecific GAT inhibitor NPA (1.5 mM) recorded in DLS (*blue*, n = 9 experiments/5 mice), NAcC (*purple*, n = 6 experiments/4 mice) or NAcS (*green*, n = 6 experiments/4 mice). Wash-on data are normalized to baseline (first 4 time points, *dotted line*) and the 4 time points are used for statistical comparisons (*shaded region*). *Right*, Mean profile of $[DA]_o$ at baseline (solid transient) and after NPA wash-on (dashed transient). DHβE (1 μM) was present throughout. (**f**) Mean peak $[DA]_o$ (± SEM) evoked by 1p following GAT inhibition with NPA in DLS, NAcC and NAcS (expressed as a % of pre-NPA baseline). Statistical significance was assessed by Kruskal-Wallis tests and Dunn's multiple comparisons. *$p$ < 0.05, **$p$ < 0.01, ***$p$ < 0.0001.

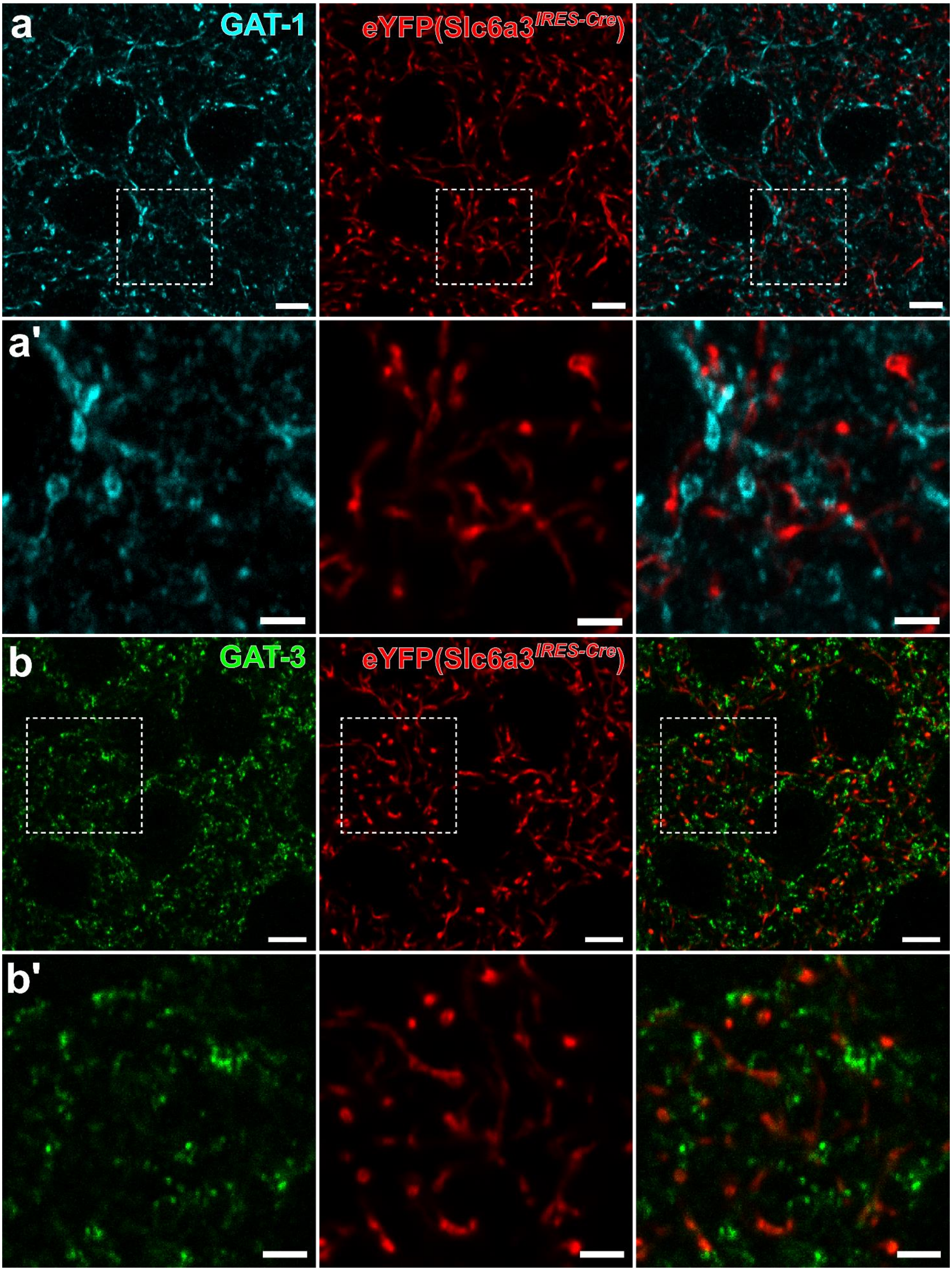


**Supplementary Fig 6. GAT-1 and GAT-3 are not typically co-localized to DA axons.**
(**a**,**b**) Representative immunofluorescence signals in dorsal striatum for GAT-1 (*cyan*, a), GAT-3 (*green*, b), and conditional eYFP expression in DA axons from Slc6a3$^{IRES\text{-}Cre}$ injected mice (*red*). (**a'**,**b'**) Immunofluorescence signals with increased magnification from a and b as indicated. We did not observe robust co-localization of GAT-1 or GAT-3 on DA axons (n = 3 mice). Scale bars: 5 µm (a,b), 2 µm (a',b').

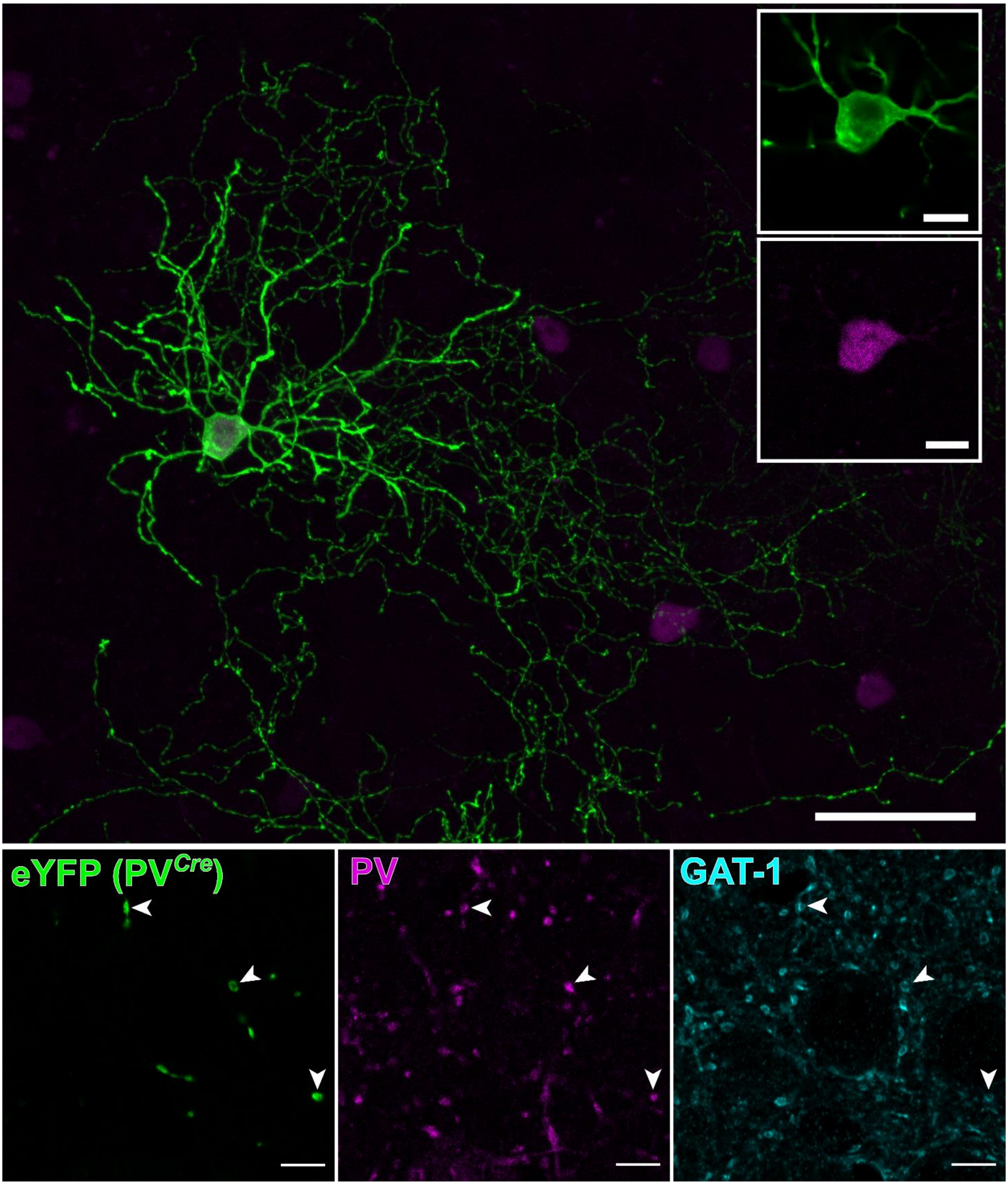


**Supplementary Fig 7. GAT-1 is co-localized to striatal parvalbumin-positive neuropil**. *Top*, Representative immunofluorescence signals for parvalbumin (PV, *purple*)-positive interneurons in the dorsal striatum, some of which also conditionally expressed eYFP (*green*) in their processes after AAV injection, in a $PV^{Cre}$ mouse. *Bottom*, We observed robust GAT-1 (cyan) colocalization in the neurites of parvalbumin/eYFP-positive interneurons (n = 3 animals). Scale bars for Top: 50 µm, 10 µm for insets. Scale bars for Bottom: 5 µm.

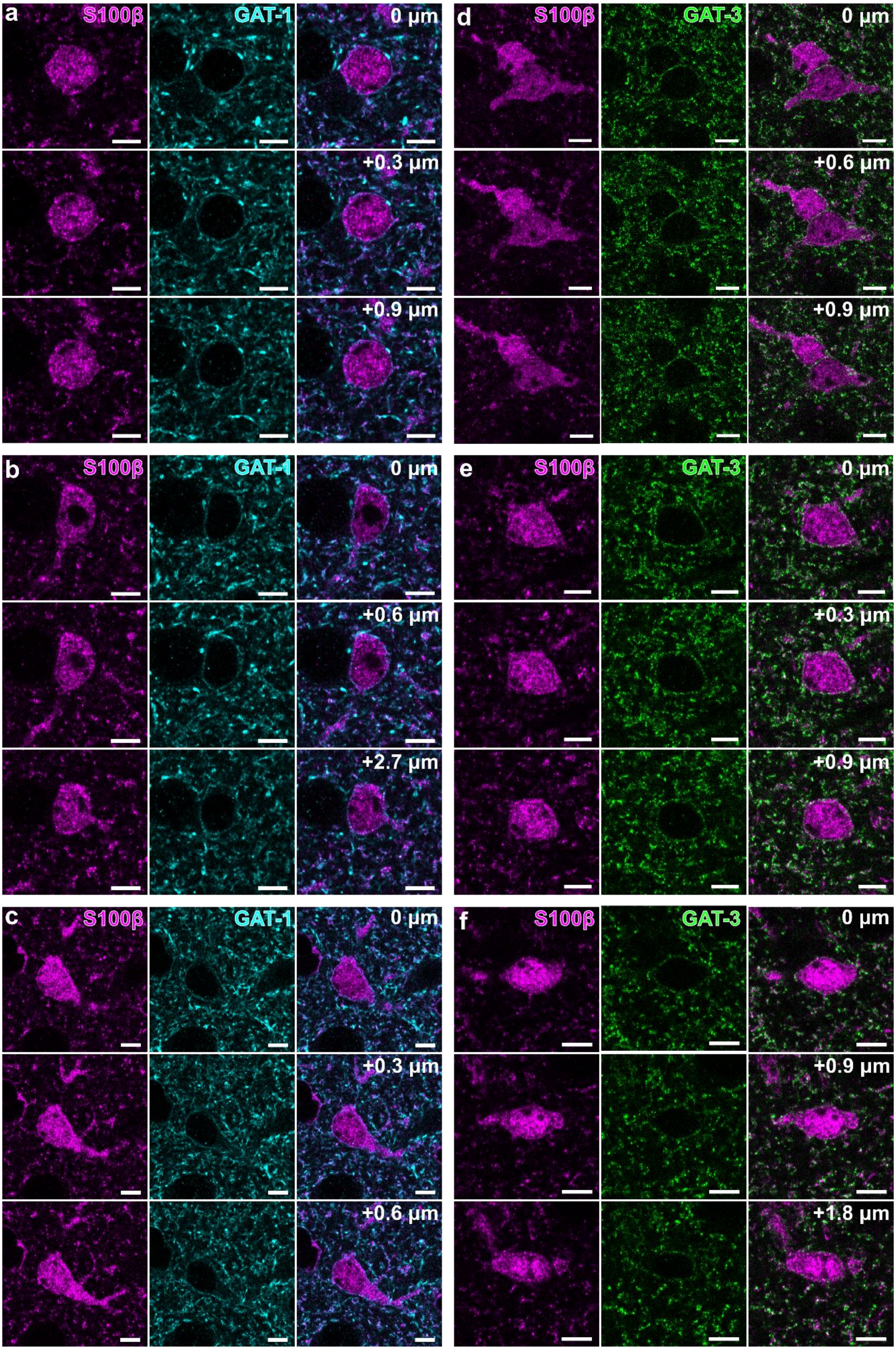


**Supplementary Fig 8. GAT-1 and GAT-3 are expressed on plasma membranes of striatal astrocytes.**
Striatal immunofluorescence signals for astrocyte marker S100β (*magenta*) and GAT-1 (*cyan*, **a**-**c**) or GAT-3 (*green*, **d**-**f**) from astrocytes represented in Fig. 5 across three separate z-planes to highlight GAT expression throughout the plasma membranes of striatal S100β-expressing astrocytes imaged in DLS. Scale bars: 5 µm.

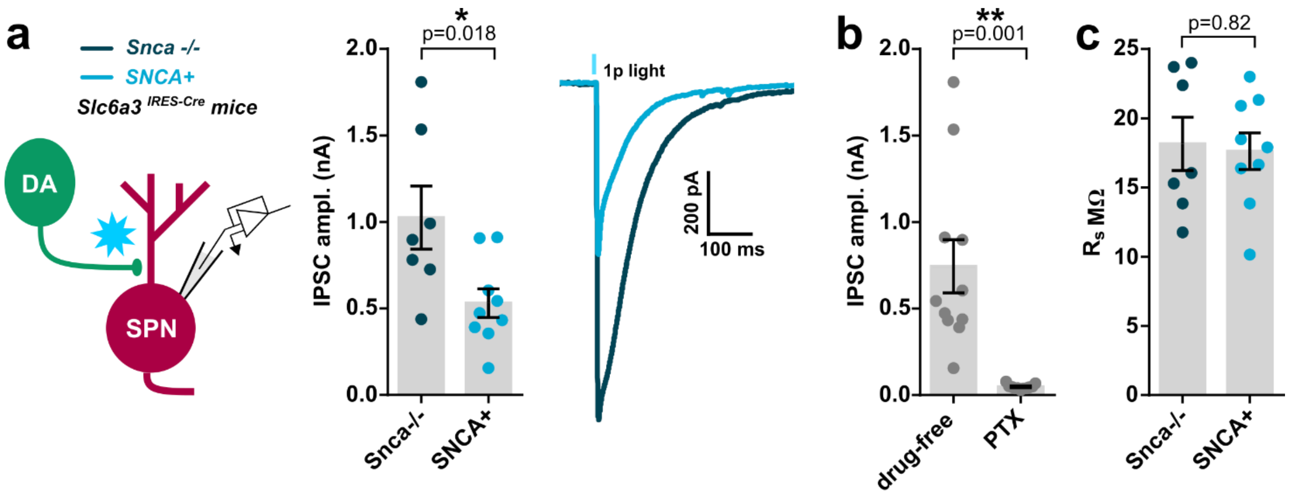


**Supplementary Fig 9. GABA co-release from DA axons in *SNCA*+ mice generates GABA$_A$ receptor-dependent IPSCs in SPNs and deficits are not due to differences in series resistance.** (**a**) Mean (± SEM) 1-pulse light evoked inhibitory postsynaptic currents (IPSCs) recorded from spiny projection neurons (SPNs) every 30s in the DLS of *SNCA*+ mice (light blue, n = 9 cells/4 mice) and littermate controls (*Snca*-/-, dark blue, n = 7 cells/4 mice), voltage clamped at -70 mV and in the presence of ionotropic glutamate receptor antagonists (NBQX, 5 µM; D-APV, 50 µM) from main Fig 8. (**b**-**c**) IPSCs in SPNs of both *SNCA*+ and *Snca*-/- mice were abolished in the presence of GABA$_A$ receptor antagonist picrotoxin (PTX, 100 µM, b) and differences in recorded IPSC amplitude not due to differences in series resistance (Rs) between recordings (c). Statistical significance assessed with Student's unpaired t-tests. *$p < 0.05$, **$p < 0.01$.
